# SRSF2 transcriptional function maintains genome integrity during cell division

**DOI:** 10.1101/2024.07.08.602469

**Authors:** Rebecca E. Wagner, Leonie Arnetzl, Thiago Britto-Borges, Anke Heit-Mondrzyk, Ali Bakr, Etienne Sollier, Jasper Panten, Sylvain Delaunay, Daniela Sohn, Peter Schmezer, Duncan T. Odom, Karin Müller-Decker, Christoph Plass, Christoph Dieterich, Pavlo Lutsik, Susanne Bornelöv, Michaela Frye

## Abstract

Recurrent mutations in serine / arginine-rich splicing factor 2 (SRSF2), particularly the proline-to-histidine substitution at position 95 (P95H), have been proposed to drive neoplastic diseases. SRSF2 plays pivotal roles in pre-mRNA processing and gene transcription. However, the precise impact of these diverse functions of SRSF2 on cancer cell behaviour remains unclear. Here, we show that deletion or homozygous P95H mutation of *SRSF2* both cause extensive DNA damage leading to cell cycle arrest *in vitro* and *in vivo,* regardless of cell genotype. We mechanistically demonstrate that SRSF2 is required for efficient bi-directional transcription of DNA replication and repair genes, independent of its function in splicing. In contrast, SRSF2 haploinsufficiency induces DNA damage without halting the cell cycle in cancer cells. Inducing the heterozygous *Srsf2* P95H mutation leads to clonal expansion of epidermal cells in mouse skin, but tumor formation is inhibited following exposure to carcinogens. To survive carcinogen treatment, cells containing the *Srsf2* P95H mutation undergo substantial transcription rewiring that restores bi-directional gene expression. Thus, our study reveals crucial roles of SRSF2 beyond splicing and underscores its importance in regulating transcription to orchestrate the cell cycle along with the DNA damage response.

## Introduction

The highly conserved serine/arginine-rich (SR) protein family is composed of 12 members in humans ^1,2^. The majority of SR proteins regulate pre-mRNA alternative splicing by increasing the recognition of weak splice sites through the splicing machinery ^3^. Depending on the location of SR protein binding in introns or exons, they act as activators or repressors in a context-dependent manner and in concert with other RNA-binding proteins ^4^. The majority of SR proteins shuttle between the nucleus and the cytoplasm. SRSF2 and SRSF5 are the only family members restricted to the nucleus ^5–9^. Beyond their primary function in splicing, SR proteins have diverse roles in regulating gene expression. The shuttling SR proteins participate in mRNA export from the nucleus to the cytoplasm, impacting mRNA stability and translation efficiency ^10–18^.

Pre-mRNA splicing begins during RNA synthesis and splicing signals can also enhance gene transcription ^19^. Several SR proteins (SRSF1, SRSF2, SRSF7) have been reported to regulate RNA polymerase II (Pol II)-mediated nascent transcription ^20–23^. The underlying molecular mechanism of how SR proteins regulate transcription is best-understood for SRSF2. To engage in active transcription, the Pol II initiation complex needs to transition into the elongation complex ^24^. This initiation-to-elongation transition requires the C-terminal domain (CTD) of Pol II to be phosphorylated by cyclin-dependent kinases (CDKs) mediating the CTD’s interaction with transcriptional regulators ^25^. During this process, SRSF2 binds to nascent RNAs to promote CDK9-activation, and thereby facilitate the release of Pol II into productive elongation ^21,26^.

Like most SR proteins, SRSF2 is essential for normal development. Germ line deletion of SRSF2 is embryonic lethal in mice ^27,28^, and its conditional inactivation triggers organ defects in thymus, heart and liver ^27,29,30^. Expression of SRSF2 is altered in about 2% of all cancers. Among these, acute myeloid leukaemia (AML), myelodysplastic syndromes, breast invasive ductal carcinoma, and chronic myelomonocytic leukaemia exhibit the highest prevalence of SRSF2 alterations ^31^. Mutations in the *SRSF2* gene are frequently found in myelodysplasia ^32^. The most common hot spot mutation is a substitution of proline for histidine at position 95 (P95H) leading to altered recognition of exonic splicing enhancers resulting in aberrantly spliced transcripts ^33,34^. In myeloid diseases, SRSF2 mutations are linked to abnormal differentiation processes of hematopoietic stem and progenitor cells (HSPCs) ^35,36^. Yet, how precisely mutated SRSF2 impairs tissue homeostasis and how this predisposes to cancer development remains unclear.

Here, we set out to identify fundamental cellular functions of SRSF2 in a stratified squamous epithelium, the interfollicular epidermis ^37^. To decipher how the functions of SRSF2 in splicing and transcription regulate cell behaviour, we depleted or mutated SRSF2 in normal epidermal and cancer cells. Loss of a functional SRSF2 caused DNA damage leading to cell cycle arrest *in vitro* and *in vivo* and independently of the cells’ genotype. The enhanced DNA damage in SRSF2-depleted cells was linked to inefficient transcription of bi-directionally orientated DNA replication and repair genes in mouse and human. Aberrant splicing did not explain the down-regulation of bi-directionally transcribed genes. Instead, SRSF2 was required for efficient RNA Pol II pause release and transcription elongation, and bi-directional genes were particularly vulnerable to loss of SRSF2. In cancer cells, the interconnection between cell cycle arrest after DNA damage was disrupted when the expression levels of SRSF2 were decreased by two-fold. SRSF2 haploinsufficiency caused DNA damage but not cell cycle arrest. Unexpectedly, inducing the SRSF2 P95H heterozygous mutation in mice did not enhance epidermal tumour formation following exposure to carcinogens. However, clonal expansion of SRSF2-mutant epidermal cells was not impaired and transcription of bi-directional genes was rescued. Thus, we show that SRSF2 serves a key role beyond splicing, and its function in regulating efficient transcription is required to coordinate the cell cycle with the DNA damage response.

## Results

### SRSF2 is required for cell cycle progression in normal epidermal and cancer cells

The functional role of SRSF2 in the mammalian epidermis is unexplored. Therefore, we depleted SRSF2 in primary human keratinocytes (HK), a well-characterised *in vitro* model for interfollicular epidermal cells ^38^. To identify functions of SRSF2 in cancer, we included two oral squamous cell carcinoma (SCC) lines, the well-differentiated SCC25 and the undifferentiated FaDu cells (Figure 1A-D; Supplementary Figure 1A). Knock-down of SRSF2 reduced cell proliferation in all three cell lines (Figure 1E; Supplementary Figure 1B, C). Cell cycle arrest was caused by a reduced ability to enter S-phase, where DNA replication occurs, leading to accumulation of SRSF2-depleted cells in sub-G0 (Figure 1F, G). The inability to efficiently initiate DNA replication in the absence of SRSF2 was confirmed by consistent down-regulation of CDC45 (Supplementary Figure 1D) ^39,40^. Normal keratinocytes undergo differentiation when exiting the cell cycle, but the cancer lines showed no increased in expression of differentiation markers KRT10 and TGM1 (Supplementary Figure 1E) ^40^. Surprisingly, cell cycle arrest only modestly induced cell death 72 hours after SRSF2-depletion (Figure 1H, I; Supplementary Figure 1F). We concluded that acute loss of SRSF2 caused cell cycle arrest likely due to impaired DNA replication.

**Figure 1.**
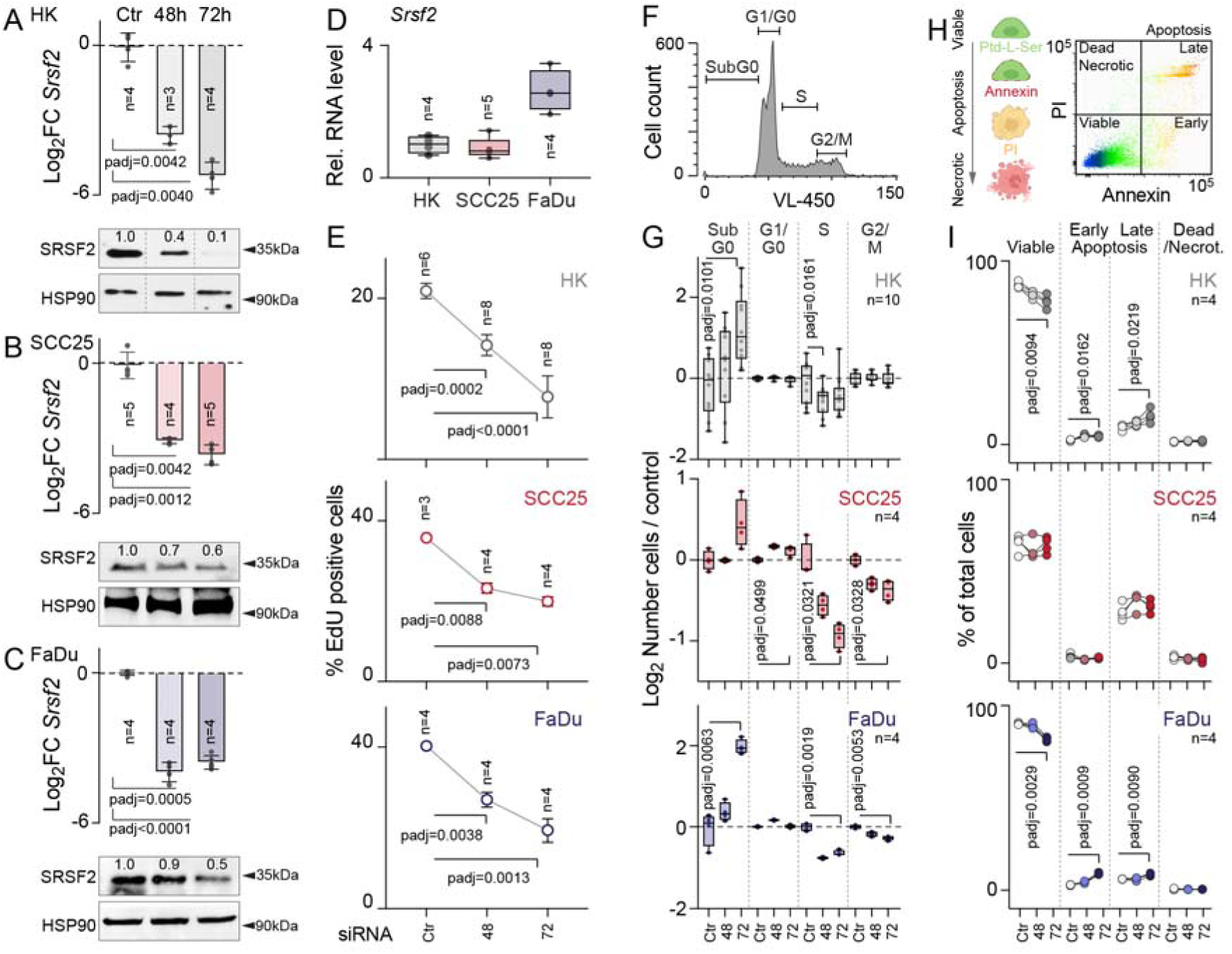
SRSF2 is required for cell cycle progression. **(A-C)** Change of *SRSF2* RNA (top) and protein (bottom) levels in primary human keratinocytes (HK) (A) and the squamous cell carcinoma cells SCC25 (B) and FaDu (C) transfected with a scramble (Ctr) for 72 hours (h) or *SRSF2* siRNA for 48h or 72h (n = number of transfections averaged over 3 technical replicates). HSP90: loading control. Numbers in Western Blot: Normalized intensity of bands relative to Ctr. (**D**) SRSF2 RNA levels in HK, SCC25, and FaDu cells (n = number of RT-QPCRs average over 3 technical replicates). (**E**) Percentage (%) of EdU-positive cells in HK (top), SCC25 (middle) and FaDu (bottom) cells transfected with Ctr for 72h or SRSF2 siRNA for 48 and 72h (n = number of transfections). (**F**) Illustration of cell cycle phase assignments. (**G**) Number of *SRSF2*-depleted HK (top), SCC25 (middle) and FaDu (bottom) cells in the indicated phases of the cell cycle 48h and 72h after transfection relative to Ctr (72h) (n = number of transfections). (**H**) Illustration of Annexin viability assay. (**I**) Percentage (%) of viable HK (top), SCC25 (middle) and FaDu (bottom) cells transfected with Ctr (72h) or *SRSF2* siRNA for 48 or 72h. (n = number of transfections). Shown is mean ± SD (A-C top panels, E,L). Box plots show minimum, first quartile, median, third quartile, and maximum (D,G). Dunnett’s multiple comparisons test (A-C top panels, D,E,G,I). Exact p-values are indicated.

### SRSF2 is required for efficient transcription of DNA replication and repair genes

Replication stress is commonly found in Myelodysplastic Syndrome patients with mutations in splicing factors including SRSF2, but the precise underlying molecular mechanism for this remains unknown ^41^. To determine how depletion of SRSF2 caused cell cycle arrest in epithelial cells, we performed transcriptome analyses in viable SRSF2-depleted cells 48 hours after transfection (Figure 2A; Supplementary Table 1). DNA replication, damage response and repair encoding genes were amongst the most mis-regulated transcripts within the 1807 genes commonly altered in all three cell lines (Figure 2B; Supplementary Figure 2A; Supplementary Table 2). Regulators of DNA replication and repair were in fact the only significantly enriched gene-set found amongst commonly down-regulated genes (Supplementary Figure 2B, C). We concluded that both normal epidermal and cancer cells showed signs of replication stress when SRSF2 was depleted.

**Figure 2.**
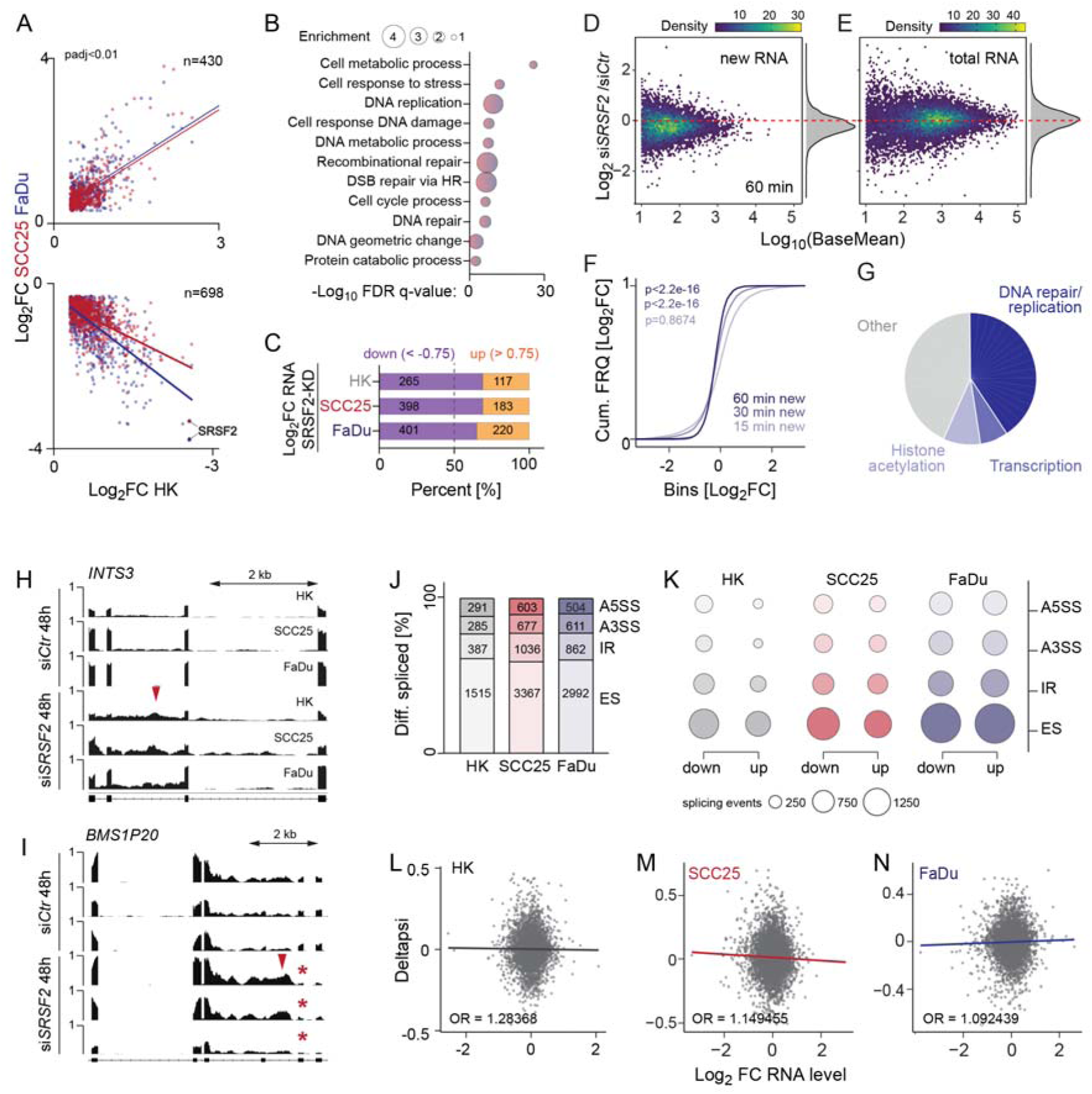
SRSF2 is required for global transcription elongation and splicing. (**A**) Correlation of shared significantly (p<0.01) up-regulated (Log_2_FC>0.3) and down-regulated (Log_2_FC<-0.3) genes between primary human keratinocytes (HK) and the cancer cell lines SCC25 and FaDu after depletion of SRSF2 for 48 hours (h) (n = differentially expressed genes). (**B**) Gene Ontology of shared significantly (p<0.01) differentially expressed genes (n=1807) in SRSF2-depleted HK, SCC25 and FaDu cells. (**C**) Percentage of down-(blue) and up-(orange) regulated genes after knockdown (KD) of SRSF2 for 48h in HK, SCC25 and FaDu cells. (**D**) Log_2_ fold-change (FC) of global nascent (new RNA) (D) and bulk (total RNA) (E) RNA levels 60 minutes (min) after 4sU metabolic labelling in si*SRSF2* versus si*Ctr* cells. Shown are all genes with a base mean >10. (**F**) Log_2_ cumulative frequency (FRQ) of nascent transcripts in si*SRSF2* versus si*Ctr* cells 15, 30 or 60 min after 4sU labelling. p= Wilcoxon rank sum test with continuity correction. (**G**) Gene ontology analyses of significantly (padj<0.05) different new transcripts 60 min after metabolic labelling. (**H,I**) Genome browser images of transcripts with intron retention (red arrowhead) leading to down-regulation (*INTS3*) or exon skipping (asterisk) (*BMS1P20*) after depletion of SRSF2. (**J,K**) Percentage of differentially (Diff.) spliced (J) and significantly (p<0.01) up-or down-regulated (K) transcripts per splicing event in SRSF2-depleted HK, SCC25 and FaDu cells after 48h. Alternative splicing was quantified using Majiq. ES: Exon skipping. IR: Intron retention. A3SS: Alternative 3’ splice site. A5SS: Alternative 5’ splice site. Total number of events are indicated in the bar charts. (**L-N**) Correlation of change in percent spliced (Deltapsi) genes with expression in *SRSF2*-depleted HK (L), SCC25 (M) and FaDu (N) cells. Deltapsi was calculated with Leafcutter. OR = odds ratio.

We also noted that across all experiments, the number of repressed genes was consistently larger than the number of up-regulated genes (Figure 2C). Therefore, we considered the possibility that SRSF2 was more globally required for efficient transcriptional activation. To directly measure whether SRSF2 regulated transcription, we quantified nascent transcription in the absence of SRSF2 by metabolically labelling RNA using 4-thiouridine (s^4^U) and performing SLAM sequencing (Supplementary Figure 2D) ^42^. Newly transcribed RNA levels significantly decreased in SRSF2-depleted cells after 30 and 60 minutes of s^4^U labelling, while total RNA levels were unaffected (Figure 2D-F; Supplementary Figure 2E-H). Almost half of the most significantly repressed nascent transcripts encoded genes regulating DNA replication and repair (Figure 2G; Supplementary Figure 2I,J; Supplementary Table 3). We concluded that SRSF2 was required for efficient gene transcription, but particularly impacted genes regulating DNA replication and repair. Therefore, we next considered the possibility that SRSF2 was required for proper splicing of genes involved in the DNA damage response.

### Aberrant splicing does not explain gene repression profiles in SRSF2-depleted cells

SRSF2 mutant cells often include poison cassette exons that introduce premature termination codons resulting in NMD ^34,43–45^. Many poison exons are essential for cell growth and frequently produce ultra-conserved RNA isoforms ^46^. We readily identified poison exon inclusions in all SRSF2-depleted cells (Figure 2H, I; red arrowheads). However, genes known to contain poison exons were not consistently repressed, and also not enriched in regulators of DNA replication and repair (Supplementary Figure 2K, L) ^46^. Therefore, we extended our analyses to all types of alternative splicing. We found similar splicing patterns in all three SRSF2-depleted cell lines (Figure 2J, K). However, a correlation between differential splicing and change in gene expression was lacking (Figure 2L-N), and the commonly altered spliced transcripts (n=221) showed no significant enrichment of any specific regulators in gene ontology analyses (Supplementary Figure 2M). We concluded that the transcriptional repression of DNA replication and repair genes in SRSF2-depleted cells was not directly caused by their mis-splicing, and considered the possibility that SRSF2 directly regulated transcription of DNA replication and repair genes.

### SRSF2 orchestrates Pol II bi-directional transcription

SRSF2 was reported to promote Pol II pause release via binding to nascent RNAs ^21,47^. Our own analyses confirmed that SRSF2 preferentially located close to transcriptional start sites, indicating a role of SRSF2 in transcriptional activation as splicing does not occur in exon 1 (Supplementary Figure 3A, B) ^26^. In the absence of SRSF2, Pol II occupancy was enhanced at transcription start sites (TSS) (Figure 3A, B; Supplementary Figure 3C, D), and the release of Pol II into active elongation was delayed (Figure 3C). Enhanced Pol II stalling and reduced transcription elongation was further confirmed by decreased Pol II phosphorylation at serine 2 (Ser2), while Ser5 phosphorylation remained unaltered (Supplementary Figure 3E, F) ^48^. In summary, our data demonstrated that loss of SRSF2 caused Pol II stalling, but why DNA replication and repair genes were most sensitive to SRSF2-depletion was still unclear.

**Figure 3.**
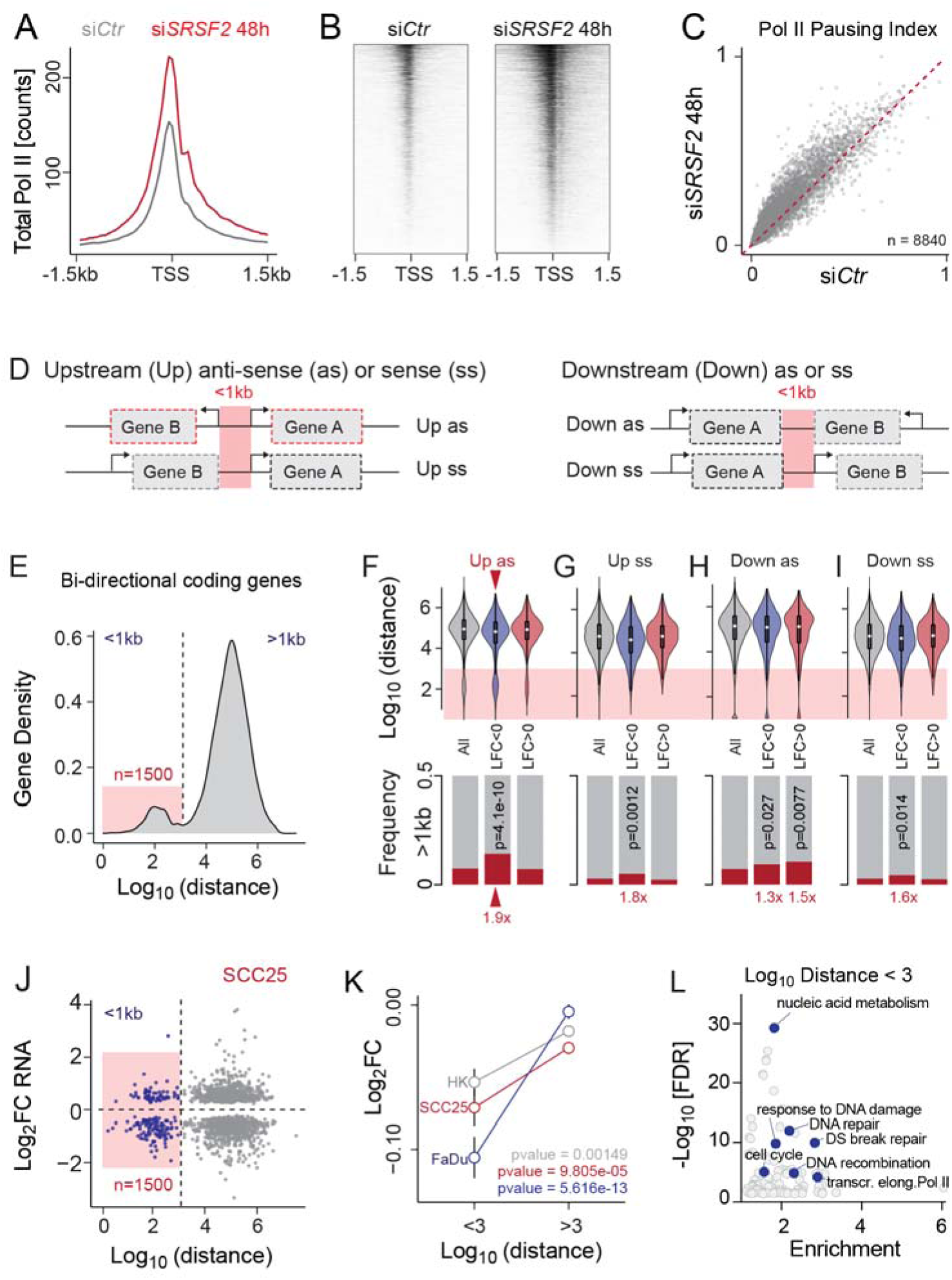
Bi-directionally transcribed gene pairs are most sensitive to SRSF2-depletion. (**A**) Genome-wide Pol II occupancy 1.5 kb up-and down-stream of transcriptional start sites (TSS) in control (si*Ctr*) and SRSF2-depleted (si*SRSF2*) FaDu cells 48 hours (h) after transfection. Shown is one out of three replicates. (**B**) Heatmaps showing total Pol II coverage at TSS shown in (A). Shown is one out of three replicates. (**C**) Correlation of pausing index (ratio of total Pol II at TSS versus the gene body) in si*Ctr* and si*SRSF2* cells (n = genes). (**D**) Illustration of possible gene orientations less than 1 kb apart from each other. Red dotted square: Bi-directional genes. (**E**) Histogram of genes separated by their distance to their closest up-stream anti-sense orientated neighbour (n = genes). (**F-I**) Distance of all (grey) or significantly (padj<0.05) up-(red; Log_2_ (L)FC > 0), or down-(blue; LFC < 0) regulated genes in SRSF2-depleted cells (upper panel) and frequency of genes less than 1 kb apart (lower panel) in up-stream (Up) antisense (as) (F) and sense (ss) (G) or down-stream (Down) as (H) or ss (I) orientation. Grey box: All genes less than 1 kb apart. Genes defined as up or down were compared to all genes, p = Fisher’s Test. (**J**) Common differentially expressed (p < 0.01) genes in SCC25 cells (n = 1807) separated by their distance to the next up-stream protein coding genes in anti-sense direction. (**K**) Log_2_ fold change (FC) of bi-directionally transcribed genes less than 1 kb (<3) or more than 1kb away (>3) from each other in HK, SCC25 and FaDu cells. P = Wilcoxon rank sum test with continuity correction. (**L**) Gene ontology analysis of 1500 bi-directional genes highlighted in red in (E).

Similar to many histone genes, genes regulating DNA replication and repair are often co-regulated through bi-directional promoters, which are DNA regions less than 1kb in length and up-stream of two adjacent genes orientated in opposite directions (Figure 3D, top left panel; red boxes) ^49–54^. The orientation of Pol II and the spatial constraint at bi-directional promoters both pose challenges to the transcription machinery. To avoid collisions, Pol II undergoes promoter-proximal pausing in the sense (ss) and anti-sense (as) directions of the respective transcriptional start sites (TSS) (Core et al., 2008, Duttke et al., 2015, Flynn et al., 2011).

We identified a distinct group of about 1500 genes having an anti-sense transcribed protein coding gene in close proximity (<1kb) in the human genome (Figure 3E; red box; Supplementary Table 4). Indeed, we found that bi-directionally transcribed coding genes were amongst the most significantly down-regulated RNAs in all SRSF2-depleted cell lines when compared to otherwise orientated genes (Figure 3F-I; red arrowhead). Bi-directionally transcribed gene pairs less than 1kb apart were significantly down-regulated in all three cell lines, while bi-directionally orientated genes with a distance greater than 1kb from each were largely unaffected by SRSF2-depletion (Figure 3J, K; Supplementary Figure 3G-I). Since SRSF2 is required for efficient transcriptional elongation, we also tested whether gene length contributed to global gene repression in the absence of SRSF2. However, the relationship between gene length and expression was uncorrelated (Supplementary Figure 3J-M). Finally, we confirmed that bi-directionally transcribed genes were indeed significantly enriched in regulators of DNA damage and repair (Figure 3L). In contrast, genes orientated in an anti-sense direction but with a distance of >1Mb frequently regulated cell adhesion and growth (Supplementary Figure 3N).

In summary, our data showed that SRSF2 was required for efficient transcriptional activation genome-wide. However, bi-directionally transcribed protein coding genes were particularly sensitive to loss of SRSF2 due to their unique organization in the genome resulting in spatial constraints and potential collision of transcriptional machineries.

### SRSF2 licences cell cycle to protect from DNA damage during divisions

To demonstrate that repression of DNA replication and repair genes resulted in increased DNA damage in SRSF2-depleted cells, we performed comet assays that detect DNA strand breaks at high resolution (Figure 4A; Supplementary Figure 4A) ^55^. Unlike normal cells, cancer cells exhibited enhanced DNA damage as early as 48 hours after SRSF2 knockdown (Figure 4B; Supplementary Figure 4B), a time point where cell viability was unaffected (Figure 1I).

**Figure 4.**
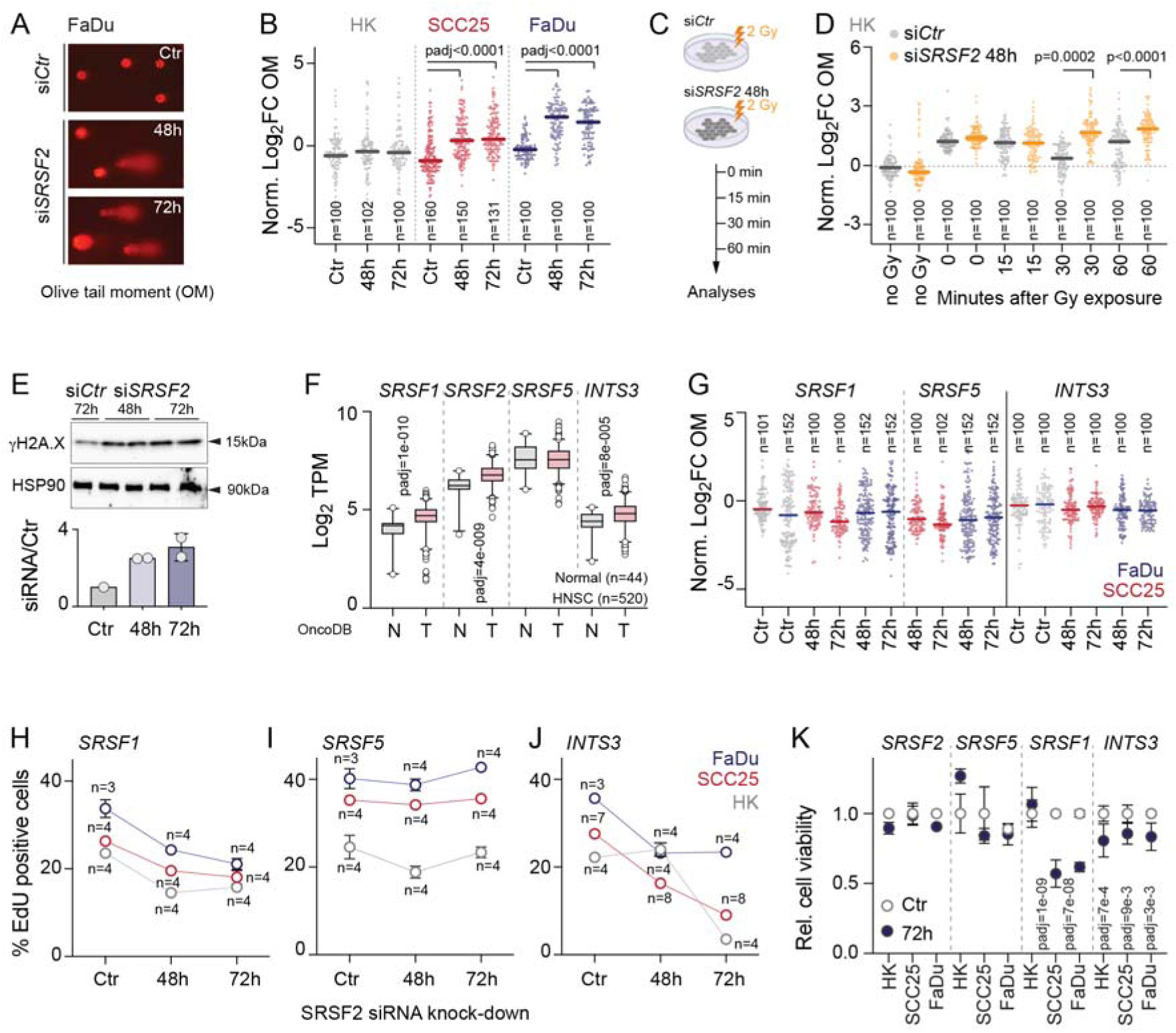
SRSF2 depletion leads to increased DNA damage without affecting cell viability. (**A**) Comet assay representative fluorescence images from control (si*Ctr*), and SRSF2-depleted (si*SRSF2*) FaDu cells 48 or 72 hours (h) after transfection. (**B**) Log_2_ fold change (FC) of the normalized (norm.) olive tail moment (OM) relative to Ctr in primary human keratinocytes (HK) and the cancer cell lines SCC25 and FaDu 48 or 72h after transfection (n = number of cells). (**C**) Experimental design for assaying DNA repair after exposure to ionizing radiation in HK cells. Gy: Gray. (**D**) Log_2_FC of norm. OM in untreated (no Gy) and treated (Gy) in HK transfected with Ctr or *SRSF2* siRNA for 48h. Values are relative to Ctr noGy (n = 100 cells averaged over two replicates). (**E**) Western Blot (upper panel) and quantification (lower panel) of γH2AX protein levels by measuring band intensity. (**F**) Log2 transcripts per million (TPM) of SRSF1, SRSF2, SRSF5 and INTS3 in normal (N) and tumour (T) samples from OncoDB (http://www.oncodb.org/). N = number of individuals. (**G**) Log_2_ FC of norm. OM relative to Ctr in HK, SCC25, and FaDu cells 48 or 72h after transfection (n = number of cells). (**H-J**) Percentage (%) of EdU-positive cells in HK (gray), SCC25 (red) and FaDu (blue) cells transfected with Ctr for 72h or SRSF1 (H), SRSF5 (I) and INTS3 (J) siRNA for 48 and 72h (n = number of transfections). (**K**) Relative cell viability of control and cells depleted for 72h with SRSF2, SRSF5, SRSF1 and INTS3 siRNAs in the indicated cell lines. Shown is mean (B,D,G) or mean± SD (H-K). Box plots show minimum, first quartile, median, third quartile, and maximum (F). Dunnett’s multiple comparisons test (K). Exact p-values are indicated.

To test whether loss of SRSF2 impaired DNA repair in response to stress in normal cells, we exposed human keratinocytes to ionizing radiation (2 Gy) and measured DNA strand breaks in a time course (Figure 4C). Cells lacking SRSF2 had significantly more DNA strand breaks 30 and 60 minutes after irradiation (Figure 4D). Up-regulation of phosphorylated γH2A.X, an early cellular marker for DNA double strand breaks, confirmed that SRSF2 was required for efficient DNA repair in all cell lines (Figure 4E; Supplementary Figure 4C). Thus, SRSF2-depleted cells accumulated DNA damage causing cell cycle arrest.

### Coupling cell cycle progression with efficient DNA repair is a unique SRSF2 function

Next, we considered the possibility that other SR proteins have similar DNA damage protective functions during the cell cycle. We tested SRSF1 and SRSF5. Similar to SRSF2, SRSF1 was reported to associate with the non-coding RNA *RN7SK*, which sequesters P-TEFb in an inhibitory small nuclear ribonucleoprotein complex ^21,56^. SRSF5 and SRSF2 are the only predominantly nuclear SR proteins ^5^. In addition, we included INTS3 into our studies because the INTS3 transcript consistently included a poison exon in the absence of SRSF2 (Figure 2H). INTS3 is a member of the integrator complex, which binds Pol II to regulate specific target genes ^57^, and aberrant INTS3 splicing by mutant SRSF2 was reported to contributed to leukemogenesis ^43^.

Similar to SRSF2, SRSF1 and INTS3 were significantly up-regulated in head and neck cancer (HNSC) tumour samples, whereas the expression of SRSF5 remained unaltered compared to normal tissue (Figure 4F). Knock-down of none of these proteins increased DNA strand breaks (Figure 4G) or specifically decreased phosphorylation of Ser2 in the Pol II CTD (Supplementary Figure 4G,H). However, phosphorylated γH2A.X was slightly induced and knocking down SRSF1, SRSF5 and INTS3 also affected cell cycle progression (Supplementary Figure 4I-M). Yet, cell proliferation was only reduced in SRSF1- and INTS3-depleted cells leading to a significant reduction of viable cells (Figure 4H-K). In summary, only depletion of SRSF2 caused cell cycle arrest and simultaneously induced stalling of Pol II leading to increased DNA damage.

### Critically low levels of SRSF2 cause DNA damage without triggering cell cycle arrest

Our finding that depletion of *SRSF2* consistently caused DNA damage and cell cycle arrest raised the question why the gene is frequently mutated in cancer ^32^. Since recurrent hotspot mutations are heterozygous, we speculated that allelic loss of *SRSF2* was sufficient to break the regulatory link between SRSF2-dependent transcription and cell cycle arrest. To test this hypothesis, we generated stable knock-down clones expressing various levels of SRSF2 using CRISPR-Cas9 gene editing (Supplementary Figure 5A-E). Mutations in *SRSF2* were highly deleterious for cancer cells. Out of 285 clones, only three carried mutations in *SRSF2* and showed a less than two-fold reduction of SRSF2 expression (Supplementary Figure 5C). The three clones (#0, #1, and #2) were used for further analyses. As a control, we generated a clone with the same CRISPR protocol but using an empty vector.

In contrast to acute deletion of SRSF2 via siRNA (Figure 1), proliferation rates were unchanged and cells with *SRSF2* gene mutations efficiently entered S-phase of the cell cycle, but then spent more time in S- and G2/M (Figure 5A, B; Supplementary Figure 5F). The extended S- and G2/M cell cycle phases can be explained by impaired DNA damage repair because SRSF2-mutant clones showed increased DNA strand breaks, a higher number of newly formed fusion genes, and up-regulated phosphorylation of γH2A.X (Figure 5C, D; Supplementary Figure 5G-I). Incorrect repair of DNA double-strand breaks can produce oncogene fusions and enhance cell survival ^58,59^.

**Figure 5.**
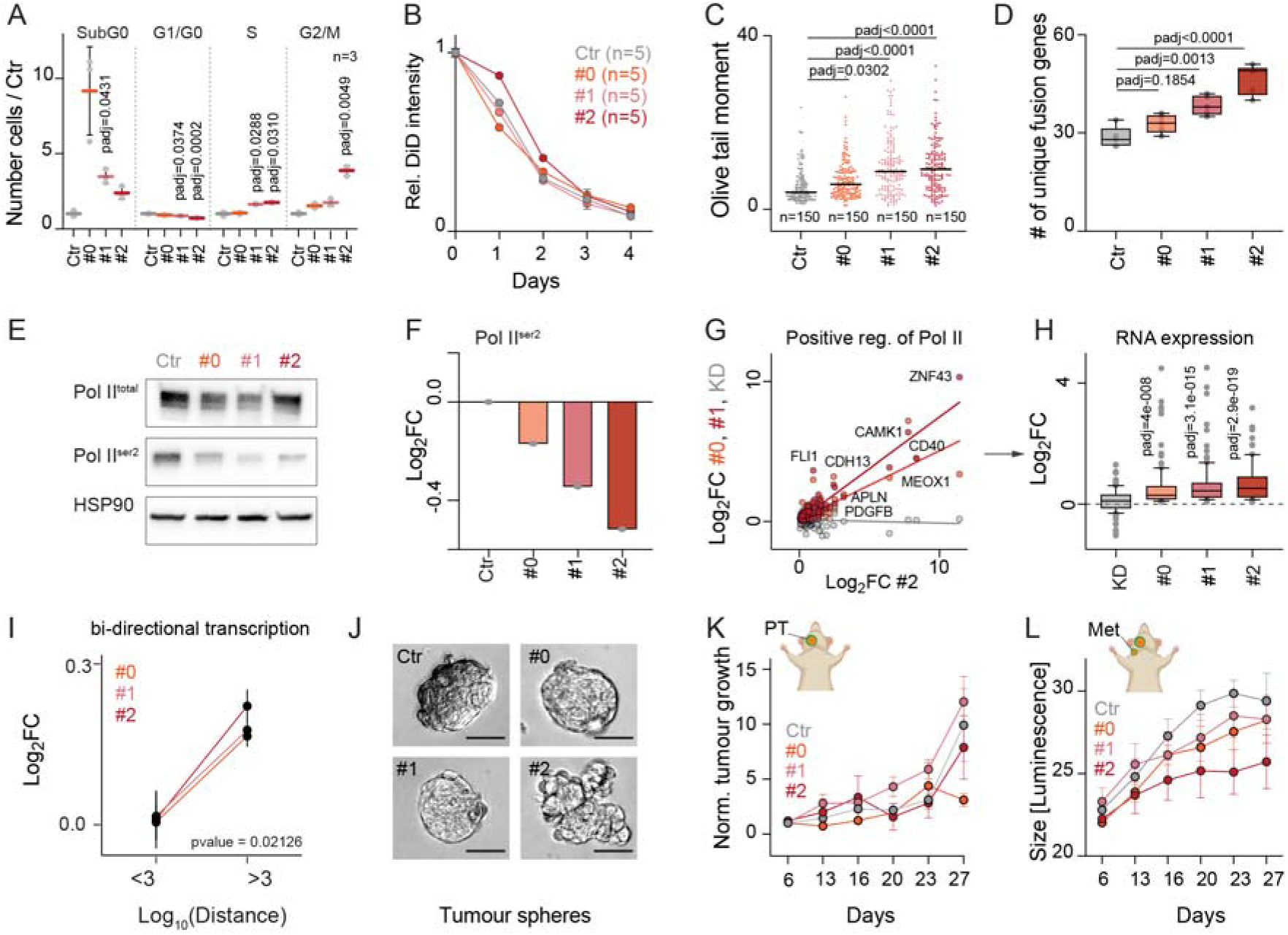
Allelic loss of SRSF2 causes DNA damage but does not impact tumorigenic potential. (**A**) Distribution of cell cycle phases in control (Ctr) or CRISPRCas9 edited clones #0, #1 and #2 repressing SRSF2 protein levels by two-fold (n = 3 flow cytometry analyses). (**B**) Proliferation rate of Ctr and #0, #1 and #2 clones relative to day 0 using DiD cell labelling. (**C**) Olive tail moment calculated from comet assay in Ctr cells and clones #0, #1 and #2 (n =150 cells). (**D**) Number of unique fusion genes in Ctr cells and clones #0, #1 and #2 (n = 5 RNA-seq experiments). (**E**, **F**) Total and serine 2 (Ser2) phosphorylated Pol II levels (E) and quantification of band intensity relative to total Pol II (Fiji) (F). (**G, H**) Correlation of Log2 fold change (FC) (G) and average (H) RNA expression levels of Pol II-positive regulators in clones #0, #1 and #2 compared to Ctr and compared to SRSF2-knockdown (KD) using siRNA for 48 hours. (**I**) Log_2_ fold-change (FC) of RNA levels of bi-directionally transcribed (<3 Log_10_ distance) compared to all other anti-sense orientated genes (>3 Log_10_ distance). (**J**) Representative images of tumour spheres cultured for 7 days formed by Ctr or #0, #1 and #2 clones. Shown is one out six experiments. (**K, L**) Normalized primary tumour (PT) growth (K) and size of lymph node metastasis (Met) (L) of orthotopically transplanted Ctr clone and clones #0, #1 and #2 into host mice monitored for 27 days. Shown is mean ± SD (A,B; J-L). Box plots show minimum, first quartile, median, third quartile, and maximum (D,H). Dunnett’s multiple comparisons test (A,C,D,H). Exact p-values are indicated.

However, regulation of transcript elongation was still impaired because phosphorylation of Pol II at Ser2 was reduced in all three *SRSF2* mutant clones (Figure 5E, F). To better understand how *SRSF2* mutant cancer cells prevented cell cycle arrest, we performed transcriptome analyses which revealed that positive regulators of Pol II were commonly up-regulated in all three clones (Figure 5G,H; Supplementary Figure 5J; Supplementary Table 5). We concluded that impaired Pol II pause release was compensated by enhancing promotor-specific transcriptional processes, because transcription of bi-directional genes was still significantly lower in all three clones (Figure 5I). Thus, the compensatory mechanisms prevented cell cycle arrest, but failed to protect from accumulation of DNA damage in SRSF2-mutated cells.

The uncoupling of cell cycle arrest and DNA damage repair did not affect the tumorigenic potential of SRSF2-mutant cells as they all grew as spheres in the absence of an extracellular matrix (Figure 5J; Supplementary Figure 5K). Moreover, the capacity of *SRSF2* mutant cells to form tumours *in vivo* was also unaltered when orthotopically injected into host mice ^60^. The number and size of primary tumours (PT) and lymph node metastases (Met) was comparable in all clones (Figure 5K, L; Supplementary Figure 5 L).

In summary, an only two-fold decrease of SRSF2 levels in cancer cells was sufficient to induce DNA damage, but this was now uncoupled from cell cycle arrest.

### The recurrent hotspot mutation P95H phenocopies Srsf2-deletion in vivo

Down-regulation of SRSF2 might not accurately recapitulate cancer-relevant functions because recurrent hotspot mutations including P95H have been reported to disrupt normal RNA splicing processes in haematological malignancies ^34,36^. However, we found that both wild-type and mutated SRSF2 proteins exhibited similar binding affinities to transcriptional start sites (Figure 6A). To test how SRSF2-deletion or -mutation affected normal epidermal cellular functions *in vivo*, we conditionally deleted or mutated (P95H) the *Srsf2* gene in the basal, undifferentiated, layer of the interfollicular epidermis in mice using an inducible oestrogen receptor domain under the control of the keratin (*Krt*) 14 promoter (*Krt14Cre-ER),* (Figure 6A-C; Supplementary Figure 6A,B). To visualize recombined cells, we included a reporter transgene (*Td-Tomato)* (Figure 6B,C, *Td-tom*; Supplementary Figure 6A).

**Figure 6.**
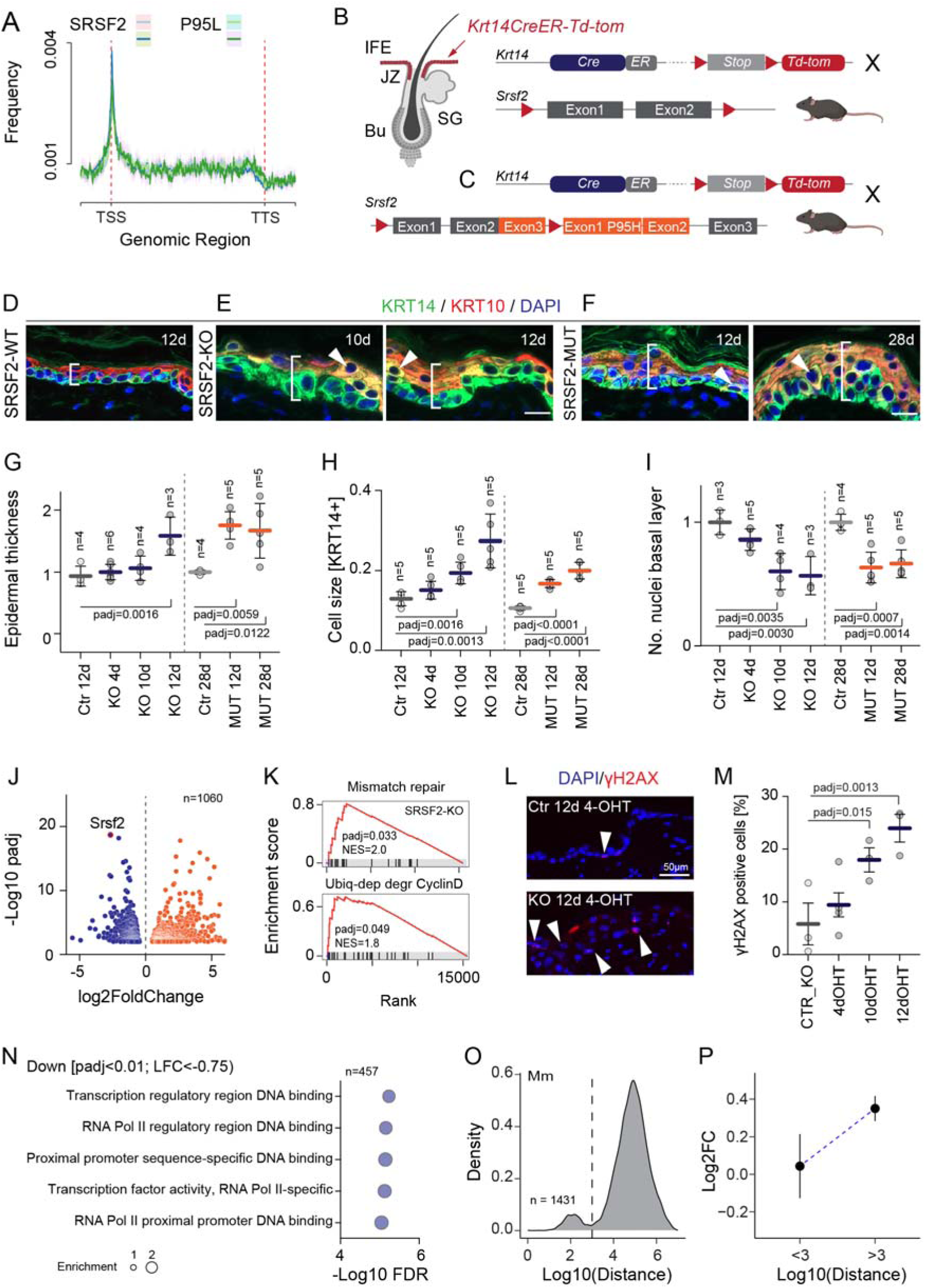
Functional SRSF2 is required for maintaining epidermal homeostasis. (**A**) Frequency of SRSF2 and mutant SRSF2 (P95L) proteins binding across genes from the transcription start site (TSS) to the transcription termination site (TTS). (**B, C**) Illustration of skin compartments (B) and transgenic lines (C). IFE: Interfollicular Epidermis; JZ: Junctional zone: Bu: Bulge; SG: Sebaceous gland. Red line and arrow: Inducible (ER) Cre-activation to delete Srsf2 in the IFE (Krt14CreER-Td-tom). Td-tom: tdTomato; Red triangles: LoxP sites. (**D-F**) Immunofluorescence for keratin (KRT) 14 and KRT10 and double-positive cells in SRSF2-WT (D), SRSF2-KO (E) and SRSF2-MUT (F) epidermis. Bracket: epidermal thickness. Arrowhead: Double-positive cells. DAPI: nuclear counterstain. Scale bars: 20 μm. (**G-I**) Quantification of epidermal thickness (G) (n = number of mice), size of cells in KRT14-positive (KRT14+) basal layer area (H) (n = 5 mice; each dot: average of 3 or 4 measurements), and total number (No.) of nuclei in basal layer (I) (n = number of mice) in SRSF2-knockout (KO) (blue) or P95H mutant (SRSF2-MUT) (orange) mice after treatment with 4-hydroxytamoxifen (OHT) for the indicated days (d). SRSF2 wildtype littermates were used as controls (Ctr_KO). Each point represents average of 9 measurement in one mouse. (**J**) Volcano plot showing differentially expressed genes in reporter-positive cells isolated from SRSF2-KO or Ctr back skin treated for 10 days (n = number of genes). (**K**) Significantly enriched gene sets in SRSF2-KO cells. NES: normalized enrichment score. (**L, M**) Immunofluorescence (L) and quantification (M) for γH2AX in SRSF2-WT and SRSF2-KO epidermis. Arrowhead: γH2AX-positive nuclei. DAPI: nuclear counterstain. (**N**) Gene Ontology (GO) analyses using significantly (p<0.01) down regulated genes (GOrilla). Background: All expressed genes. (**O**) Histogram of genes separated by their distance to their closest up-stream antisense orientated neighbour (n = genes). (**P**) Log2 fold change (FC) of all bi-directionally transcribed genes less than 1kb away (<3) or more than 1kb away (>3) in cells sorted from mouse epidermis. Shown is mean ± SD (G-I, M) or SEM (P). Dunnett’s multiple comparisons test (G-I, M). Exact p-values are indicated.

Deletion or mutation of SRSF2 both significantly increased epidermal thickness without inducing cellular proliferation (Figure 6D-F, brackets; Figure 6G; Supplementary Figure 6C-E). Instead, undifferentiated cells increased about two-fold in size leading to decreased cellularity in the basal layer of the epidermis (Figure 6H, I). As a consequence, cellular differentiation was impaired and SRSF2-depleted epidermis contained a significantly larger number of cells simultaneously expressing undifferentiation (KRT14) and early differentiation (KRT10) markers (Figure 6D-F, arrowheads; Supplementary Figure 6F-H; arrowheads). Such hybrid populations represent proliferating progenitor cells committed to differentiation ^61–63^.

In summary, we reveal that the recurrent hotspot mutation P95H phenocopies *Srsf2*-deletion and SRSF2 is required to maintain tissue homeostasis.

### Loss of Srsf2 impairs Pol II activity leading to DNA damage and cell cycle arrest in vivo

To identify why skin homeostasis was impaired in mice, we transcriptionally profiled flow sorted *Srsf2*-depleted cells (Figure 6J; Supplementary Table 6). The significant enrichment of genes involved in ubiquitin-dependent degradation of Cyclin D *Srsf2*-depleted cells indicated inhibition of G1-S phase transition ^64^. A prolonged S-phase of the cell cycle also explained the enrichment of miss-match repair genes (Figure 6K) ^65^. However, the number of DNA double strand breaks was significantly increased in cells lacking SRSF2 (Figure 6L,M).

When we inspected the down-regulated genes (n=457), we only found regulators of Pol II transcription activity to be significantly enriched (Figure 6N; Supplementary Table 7). As described for human genes (Figure 3E-L), bi-directionally transcribed mouse genes were mostly involved in regulating DNA replication and repair and on average lower expressed in the absence of SRSF2 (Figure 6O,P; Supplementary Figure 6I; Supplementary Table 8). When we compared gene expression profiles of *Srsf2*-depleted with P95H mutated epidermis, we identified 1673 commonly de-regulated genes (Supplementary Figure 6J,K). As expected, regulators of DNA replication and repair were amongst the most enriched differentially expressed genes in both mouse models (Supplementary Figure 6L).

In summary, the function of SRSF2 to promote Pol II transcription efficiency of bi-directional genes was maintained *in vivo* and evolutionary conserved in mouse and human. This is notable because splicing differences caused by the same mutation in splicing factors induce largely non-overlapping splicing changes in mouse versus human ^66^.

### SRSF2 P95H heterozygous mutation is not sufficient to drive tumour development

Finally, we asked whether induction of cell proliferation in the presence of genomic mutations was sufficient to trigger tumour development in the mouse epidermis carrying the heterozygous *Srsf2* P95H hotspot mutation. We performed classical two-stage chemical skin carcinogenesis experiments, in which tumour-initiation is achieved by induction of DNA mutations using a single dose of 7,12-dimethylbenz[a]-anthracene (DMBA), followed by stimulated epidermal proliferation and inflammation via repeated treatment with tetradecanoyl-phorbol acetate (TPA) (Figure 7A) ^67^. Under this protocol wild-type mice develop benign tumours (paillomas) at around week 10 of the treatment regimen. We induced the P95H mutations in cells from different epidermal lineages including long-lived hair follicle stem and progenitor cells (Figure 7B) ^68,69^.

**Figure 7.**
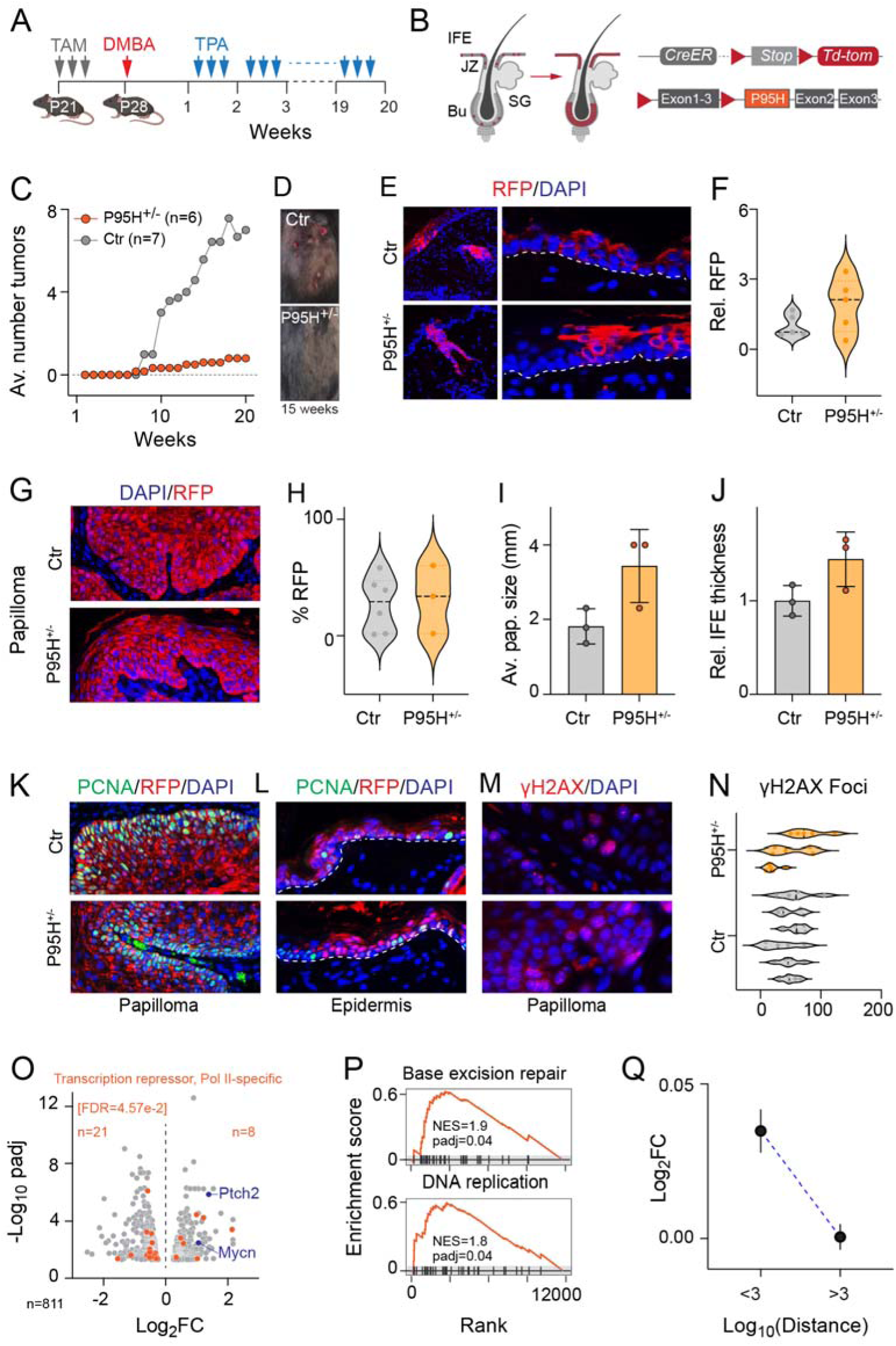
Srsf2 P95H mutation is not sufficient to drive tumour development. (**A, B**) Illustration of chemical carcinogenesis treatment regimen (A) and location of P95H cells in skin and transgenes (B). P: Postnatal day. (**C, D**) Average (av.) number of tumours (C) and examples of tumours per mouse formed over the course of 20 weeks (n = mice). (**E, F**) Control (Ctr) and P95H heterozygous (P95H+/−) epidermal cells visualized by labelling for td-Tomato (RFP) (E) and quantification of RFP-positive populations (F). (**G-I**) RFP-positive cells in papillomas (G) and quantification of percentage (H) and av. papilloma (pap.) size (I) in papillomas. Each point represents one biological replicate averaged of five images (F,H). (**J**) Epidermal thickness of Ctr and P95H+/− epidermis after the carcinogen protocol. (**K-M**) Immunofluorescence labelling for PCNA (K,L) and γH2AX (M) in papillomas (K,M) or epidermis (L) in Ctr and P95H+/− mice after carcinogenic treatment. (**N**) Quantification of (M). Dotted line: Basement membrane (E,L). DAPI: nuclear counter stain (E,G,K-M). (**O**) Volcano plot showing significantly (padj<0.01) de-regulated genes in P95H+/− compared to ctr tdTomato-positive cells. Orange: Genes of the GO: 0001227. Blue: Mycn and Ptch2. (**P**) Significantly enriched gene sets in SRSF2-P95H+/− cells. NES: normalized enrichment score. (**Q**) Log_2_ fold change (FC) RNA levels of all expressed bi-directionally transcribed (<3) and all other (>3) genes. Shown is average (C) or mean ± SD (I,J). Violine plot shows average with median (black dotted line) (F,H).

The number of tumours in mice carrying the P95H heterozygous mutation was dramatically reduced (Figure 7C,D; Supplementary Figure 7A). Unexpectedly, the P95H mutation did not affect clonal expansion of epidermal cells, and clonal growth was even slightly enhanced in P95H+/− mice (Figure 7E,F). Irrespective of genotype, one of three papillomas contained recombined tumour cells (Figure 7G,H). The size of the papillomas and the thickness of the IFE were both slightly increased in P95H+/− mice and accordingly, we found no difference in the number of cycling PCNA-positive cells (Figure 7I-L). The number of γH2A.X-positive cells was also comparable between control and P95H+/− mice (Figure 7 M,N). Thus, the heterozygous P95H mutation inhibited tumour formation following carcinogen treatment, yet this was not due to reduction in cell cycle or enhanced DNA damage.

To determine how P95H epidermal cells differed from wild-type clones, we transcriptionally profiled recombined cells after sorting for td-Tomato (Supplementary Table 9). We noted that genes regulating Pol II-specific transcription were still de-regulated in P95H clones, but now the most significant gene ontology term comprised of DNA-binding transcription factors decreasing transcription of specific genes transcribed by Pol II (GO: 0001227) (Figure 7O; Supplementary Figure 7B). The expression of the majority of these Pol II repressors was indeed decreased in P95H epidermal cells (Figure 7O; orange dots). Among significantly up-regulated genes were *N-Myc* and *Ptch2* (Figure 7O; blue dots), dysregulation of both of these genes commonly drives basal cell carcinomas ^70^.

However, P95H+/− epidermal cells only survived the carcinogenesis protocol when the transcriptional deficiency at bi-directional promoters in SRSF2 mutant cells was rescued and accordingly, expression of both DNA replication and repair genes was enhanced (Figure 7P,Q). These viable and cycling P95H mutant epidermal cells were metabolically different from wild-type cells and showed significant repression of interleukin signalling (Supplementary Figure 7C), indicating that inflammation was not induced by TPA in SRSF2 mutant cells. TPA is a protein kinase C activator, which normally induces inflammation and epidermal hyperplasia, after application to the skin ^71^. A lack of the pro-inflammatory response may explain the reduced tumour formation in the epidermis of P95H+/− mice ^72^.

In summary, exposure to carcinogens led to the emergence of viable, dividing P95H mutant cells that differed dramatically from wild-type epidermal cells including a rescue from inefficient Pol II-dependent transcription of DNA replication and repair genes.

## Discussion

SRSF2 serves dual roles as a splicing factor and transcriptional activator ^73^. The interdependence of these two processes poses a challenge when deciphering the precise underlying molecular mechanisms by which SRSF2 operates. Nevertheless, our study revealed that SRSF2 was required for efficient transcription globally, and in particular of bi-directional DNA replication and DNA repair genes. Enhancing the transcription of DNA replication and repair genes is a splicing-independent function of SRSF2. We showed that SRSF2 regulated transcription independently of cell genotype, and this function was conserved in human and mouse. In stark contrast, alternative splicing varies considerably between cell types, tissues, and organisms ^74–76^. Accordingly, we observed a very limited number of shared alternative splicing events between SRSF2-depleted cell lines, and the small group of consistently spliced genes was not enriched in specific cellular functions or processes. However, we cannot exclude that alternative splicing contributed to the observed cellular phenotypes.

The high frequency of *SRSF2* mutations in neoplastic diseases suggested that these mutations drive oncogenesis ^77^. In line with this hypothesis, mutations in *SRSF2* are among the most common in clonally expanded blood cells ^78,79^. However, we identify *SRSF2*-P95H as a loss-of-function mutation, causing extensive DNA damage leading to cell cycle arrest *in vitro* and *in vivo*. Cell cycle arrest in the absence of a functional SRSF2 is at odds with the finding that P95H alone is sufficient to induce dysplastic features and transformation into AML-like blasts ^80^. However, we also observed a higher level of double strand breaks in the absence of SRSF2. Co-ordinating DNA repair and cell cycle progression is required for cellular homeostasis. Mutations occur as a result of mutagens, or endogenous genotoxic stresses such as transcription, replication and oxidative phosphorylation, but they only accumulate when DNA repair pathways are inactivated or saturated ^81,82^. Higher rates of mutations are the source of genome instability, and increased genetic diversity can drive tumorigenesis through accelerated clonal evolution ^82,83^.

We uncovered that mutating the *SRSF2* gene is highly deleterious for cells. Only three out of 285 CRISPR/Cas9 edited clones overcame the cell cycle block and survived. Homozygous mutations in the coding regions of SRSF2 are lethal as only one heterozygous clone survived. Homozygous mutations were only tolerated in the 3’ UTR. In all cases, SRSF2 expression was down-regulated by up to 50%. All surviving clones exhibited reduced levels of Pol II phosphorylation the Serine 2 required for transcription elongation, underscoring an essential role or SRSF2 in regulating CDK9 activity. The surviving SRSF2-mutant clones compensated lower transcription levels by up-regulating positive regulators of Pol II to allow proliferation *in vitro* and *in vivo*. Nevertheless, all clones retained elevated levels of DNA damage, providing further evidence that SRSF2 is indispensable for efficient transcription of DNA repair genes.

Clonal haematopoiesis in humans is caused by the accumulation of somatic mutations over time. The *SRSF2*-P95H mutation is found in the most rapidly expanding hematopoietic clones, indicating that it drives clonal expansion ^84^. However, we provide evidence that the *SRSF2*-P95H mutation alone is not sufficient to drive oncogenic transformation. The heterozygous induction of the mutation in epidermal cells exposed to carcinogens inhibited skin tumour formation in mice. The P95H mutant epidermal cells did however clonally expand, and even showed slightly enhanced growth competition. Notably, all surviving cell clones had rescued the down-regulation of bi-directionally transcribed genes.

In summary, we reveal that SRSF2’s function as a transcriptional activator maintains genome stability in coordination with cell cycle progression. Loss of SRSF2 caused Pol II stalling and global repression of transcription and particularly strongly suppressed bi-directionally transcribed genes, which are commonly involved in DNA replication and repair. In cases where cells overcome cycle arrest through oncogenic mutations for example, loss of SRSF2 might accelerate clonal evolution by increasing DNA damage. Our findings raise the possibility of a SRSF2-driven common function in transcription supplementary to or independent of splicing contributing to the aetiology of myelodysplastic syndromes.

## Material and Methods

### Mice

Mice were housed in the central animal Laboratory at the DKFZ. All mouse husbandry and experiments were carried out in accordance with guidelines of the local ethics committee under the terms and conditions of the animal licences G-215/19, G-351/19, and G-96/21.

### Generation of the transgenic lines

Rosa-CAG-LSL-*tdTomato* (007905, The Jackson Laboratory) ^85^, *Krt14*-cre/ERT (005107, The Jackson Laboratory) ^86^, *Srsf2*^fl/fl^ (018019, The Jackson Laboratory) ^87^ and *Srsf2*^P95H^ (028376, The Jackson Laboratory) ^34^ mice have been described previously.

To conditionally delete *Srsf2* in the basal undifferentiated layer of the epidermis, *Srsf2*^fl/fl^ mice were crossed with *Krt14*-CreERT and Rosa-CAG-LSL-*tdTomato* mice. The P95H mutation was induced in the epidermis by crossing *Srsf2*^P95H^ mice with *Krt14*-CreERT mice. To activate CreERT the back skin of mice was shaved and then topically treated with 7.14 mg/ml 4-hydroxytamoxifen (Sigma Aldrich) solution in acetone every other day for the indicated period of time.

To induce the *Srsf2* P95H mutation in proliferative epidermal compartments of the skin, *Srsf2*^P95H^ mice were crossed with Krt14-CreERT and/or Lgr5-CreERT and Rosa-CAG-LSL-tdTomato mice.

### Orthotopic transplantation assays

*SRSF2* mutant clones were transduced with a lentiviral vector expressing luciferase-mCherry. Top 25% of cells expressing highest mCherry levels were flow sorted and used for transplantation assays. A total of 100,000 cells were injected into the tongue of NSG female mice. The tumour growth was monitored using *in vivo* imaging (Xenogen IVIS Imaging System-100; Caliper Life Sciences) once in the first week and then 2-3 times per week for a maximum of 4 weeks. In order to visualise the tumour cells mice received an intraperitoneal injection of 50 μl luciferin (Promega; 5mg/ml in PBS). Isoflurane gas was used to anaesthetise mice during imaging. Data were quantified with Live Image Software v4.4 (Caliper Life Sciences).

Animals were sacrificed before this time point when the tumour reached a maximal diameter as licenced. None of the animals in this study reached this endpoint. Experiments were also stopped immediately when animals showed a hunched posture or weight loss of 20% of the initial weight. Mice with tumours were also killed if they showed signs of necrosis or inflammation associated with tumour growth. Animals with moderate dyspnoea due to metastases in the lungs were killed. Mice with signs of infection, non-healing, bloody or oozing wounds were also culled. These limits were not exceeded in any of the experiments. For each experiment, mice were killed at the same time, once one experimental group reached the humane endpoint.

### Chemical carcinogenesis

Female mice between 5 and 7 weeks old were treated with three intraperitoneal (IP) injections of tamoxifen (Sigma), if not stated otherwise. One week later, mice were shaved and once treated with DMBA (400 nMol / 200 μl acetone) (Sigma). The following week promotion with TPA (10 nMol / 100 μl acetone) (Sigma) began three times a week for up to 20 weeks.

### Tissue processing and staining

Mouse back skin was fixed in paraformaldehyde overnight in 4% PFA (Santa Cruz) and transferred to ethanol. Samples were then embedded in paraffin. 5 μm thick sections were cut using a microtome. Prior to staining, the samples were deparaffinised in xylene rehydrated in a ethanol gradient. Antigen retrieval was carried out by boiling samples in sodium citrate solution for 20 minutes. Blocking and permeabilization was done by incubating samples with 10% FBS in PBS tween-20. Antibodies were diluted in blocking solution and incubated at 4°C overnight. After washing in PBS samples were incubated with secondary antibody and DAPI in PBS for 1 hour. Samples were then washed again before mounting. The following antibodies were used: KRT14 (1:1000, Abcam Ab7800), KRT10 (1:100, Abcam Ab76318), KRT6 (1:200, Abcam Ab18586), Loricrin (1:700, Biolegend 905103), RFP (1:1000, Rockland 600-4-1-379), KI67 (1:400, Cell Signaling Technologies 9129), PCNA (1:400, Abcam Ab29), γH2AX (1:100, Cell Signaling Technologies 9718).

### Cell culture

The squamous cell carcinoma cell lines SCC25 (ATCC CRL-168TM) and FaDu (ATCC HTB-43) were obtained from ATCC (https://www.lgcstandards-atcc.org). Juvenile primary human keratinocytes (HK) (C-12001, #4382014, Promocell) or (ZHC-1116, Cellworks UK). HK were cultured in basal medium (Promocell) or KGM-gold (Lonza). FaDu cells were grown in Eagles minimum essential medium (ATCC) supplemented with 10% FBS (ThermoFisher). SCC25 cells were grown in Gibco DMEM/F12, HEPES (Life Technologies) medium supplemented with 10% FBS. All cells were grown in the presence of penicillin streptomycin (ThermoFisher) in a humidified incubator at 37°C with 5% CO_2_.

### Transfection and CRISPR Cas9 gene editing

SRSF2, SRSF1, SRSF5 and INTS3 siRNA pools (30 optimally-designed siRNAs) were obtained from siToolsBiotech. siRNAs were transfected into cells using Lipofectamine RNAiMax (ThermoFisher) according to the manufacturer’s instruction. Except, FaDu cells were transfected with double the recommended concentration of lipofectamine and siRNA. Transfection of SCC25 cells was performed using Opti-MEM (ThermoFisher) according to the manufacturer’s instruction.

Guide RNAs were ordered as DNA oligos from Sigma Aldrich (guide2: GGGTATGACCTCCCTCAAGG; guide4: TCGTTCGCTTTCACGACAAG; guide5: TCGGCAAGCAGTGTAAACGG) and cloned into the pSpCas9(BB)-2A-GFP (Plasmid #48138; Addgene). FaDu cells were transfected with 500 ng of plasmid according to manufacturer’s instructions (ThermoFisher) and cells were single cell sorted 48 hours after transfection.

### Tumoroid Assay

5000 cells were resuspended in K-FSM medium (ThermoFisher) and placed in ultra-low adherent culture dishes (Stem Cell Technologies), 6 wells per condition. The number of tumoroids per well was counted on day 3, 5 and 7 and at least 15 representative pictures were taken per condition.

### Alkaline Comet Assay

2.5 × 10^5^ cells were collected in 250 μl suspension. 50 μl of cell suspension was mixed with 350 μl 0.7% agarose (Biozym). This mixture was spread on comet slides (R&D systems) and left on ice, until the agarose solidified. Cells were then lysed at 4°C overnight in lysis buffer: 2,5 M NaCl) (Sigma), 10 mM Trizma Base (Sigma), 100 mM EDTA disodium (Biotechnik GmbH) and 1% sodium N-Laurylsarcosin (Sigma,) in distilled water. Then the pH-value was adjusted to 10 using 1 M NaOH (ROTH). 1% Triton X100 (Sigma) and 10% DMSO (ROTH) were added fresh before every experiment. Slides were then placed in an electrophoresis chamber and covered with buffe (24g NaOH and 0.744 g disodium-ETDA in 2 litres distilled water) and left for 20 minutes. Electrophoresis was performed for 20 minutes at 25 volts and 300mA. Slides were then fixed with Ethanol. DNA staining was performed using SYBR-green (ThermoFisher) diluted 1:10000 in TE buffer. Comet tail moment was measured using fluorescence microscopy at 400 x magnification. Approximately 200 cells were counted per slide.

### Immunofluorescence and immunohistochemistry

Glass cover slips were coated with Poly-D Lysine hydrobromide (Sigma Aldrich). Cells were seeded on coated coverslips, fixed with 4% PFA (Santa Cruz), and washed with PBS. Cells were permeabilized with 0.1% triton and blocking was performed with 10% FBS. Primary antibodies were diluted in 1%FBS, 0.1%tween and PBS, and cells were incubated with this mix overnight at 4°C. Cells were then washed with PBS and incubated with secondary antibody and DAPI in 1%FBS for 1 hour at room temperature. Washing was then carried out prior to mounting. The following primary antibodies were used: γH2AX (1:100, Cell Signaling Technologies), S9.6 (1:200, Kerafast).

### Western blotting

Proteins were extracted from all cell lines using RIPA buffer (50 mM Tris HCl pH 8, 150 mM NaCl, 1% Nonidet P40, 0.5% sodium deoxycholate, 0.1% SDS) supplemented with cOMPLETE protease inhibitor (Roche) and PhosphoSTOP (Sigma Aldrich). Protein extracts were separated on polyacrylamide gels and blotted on nitrocellulose membranes. Membranes were blocked for 1 hour in 10% milk/BSA in TBST and incubated overnight at 4°C with the primary antibodies. Membranes were then washed in TBST and incubated for 1 hour at room temperature with HRP-conjugated secondary antibody (1:10000 in PBS; GE Healthcare/Biocat) Membranes were washed three times in TBST before bands were visualised using ECL (GE Healthcare), using the Chemidoc gel imaging system (Biorad). The following antibodies were used: H2AXγ (1:1000, Cell Signaling Technologies), SRSF2 (1:1000, Abcam), HSP90 (1:1000, Santa-Cruz), total Pol II (1:1000, Cell Signaling Technologies), Pol II ser2 (1:1000, Abcam), Pol2 ser5 (1:1000, Abcam), SRSF1 (1:1000, Cell Signalling Technologies), SRSF5 (1:1000, Abcam), INTS3 (1:1000, Bethyl Laboratories), GAPDH (1:1000, Santa Cruz).

### RNA extraction and qPCR

RNA was extracted from cells and tissues using TRIZOL (ThermoFisher) following manufacturer’s instructions and quantified using Nanodrop (NanoDrop One; ThermoFisher). cDNA was synthesised from 1 μg of RNA using Superscript III (ThermoFisher) following manufacturer’s instructions using random primers (Promega). RT-qPCRs were carried out on a Quantstudio 3 Applied Biosystems real-time PCR system (ThermoFisher) using either fast SYBR green master mix (ThermoFisher) or Taqman Fast Universal Master Mix (ThermoFisher). The following probes were used to amplify selected genes: Cdc45 (Hs00907337_m1), Cdc25 (Hs00156411_m1), Cdk1 (Hs00938777_m1), Mcm10 (Hs00218560_m1), Itga6 (Hs01041013_m1), Krt10 (Hs00166289_m1), Tgm1 (Hs00165929). The following primers were used: Srsf2_fwd:CCTAATTTGTGGCCTCCTGA, Srsf2 Rvs: TCAATCTCTTGACAGCTTTAGGC. Results were normalised to 18S rRNA (Fwd: GTAACCCGTTGAACCCCATT, Rvs: CCATCCAATCGGTAGTAGCG) (Hs99999901_s1).

### Cell cycle profiling

All cells were collected with trypsin and fixed in ice-cold 70% ethanol at least overnight. Cells were pelleted and resuspended in PBS with DAPI (1:3000) and analysed with a BD FACS Canto (BD Biosciences) analyser. Data Analysis was performed using FlowJo software. G0/G1, S, G2/M and SubG0 peaks were manually delimited on the 405 nm histogram. Statistical analysis was performed in GraphPad Prism.

### Click it 5-ethynyl-2’-deoxyuridine (EdU) and membrane labelling (DiD)

The proportion of cells in S phase of the cell cycle was assayed via EdU incorporation. Briefly, transfected or genome edited cells were incubated with EdU at a final concentration of 10 μM for 1 hour at 37°C. After 1 hour, cells were pelleted and fixed in 4% PFA (Santa Cruz). Permeabilization was performed by incubating cells with ice cold permeabilization buffer (PBS, 3% FBS and 0.1% Saponin) for 5 minutes at room temperature. Cells were then washed and click-it chemistry was carried out using Click-iT Cell Reaction Buffer kit (ThermoFisher) according to manufacturer’s instructions. Samples were analysed on a BD FACS Canto (BD Biosciences) analyser.

Cell membranes were labelled using DiD (ThermoFisher) according to the manufacturer’s instructions. In brief, cells were trypsinised and resuspended at a concentration of 1×106/ml in serum free EMEM (ATCC). Cells were then incubated at 37°C for 20 minutes. They were then centrifuged at 1500 rpm for 5 minutes at room temperature. The supernatant was washed with culture medium containing 10% FBS and pen/strep a total of three times. Cells were allowed to recover at 37oC for a minimum of 10 minutes before analysing on a BD FACS Canto. Measurements were taken every 24 hours over 5 days.

### Cell death assay

Cell death was measured by FACS analysis using propidium iodide (PI) and Annexin V staining (BD Biosciences). In brief, the supernatant and trypsinized cells were collected and centrifuged for 5 minutes at 1,800 rpm at 4 °C. Pellets were resuspended in binding buffer containing 4.3 μg/ml propidium iodide and Annexin V for 15 minutes at room temperature in the dark. Cells were analysed by FACS within 1 hour after staining. Cells were labelled as follow: live cells are PI- and AnnexinV-, early apoptosis cells are PI- and AnnexinV+, late apoptosis cells are PI+ and AnnexinV+, dead or necrotic cells are PI+ and AnnexinV-. Samples were analysed on a BD FACS Canto (BD Biosciences) analyser.

### RNA library generation

For RNA sequencing experiments, RNA was extracted from cells or tissues using TRIZOL (ThermoFisher) following manufacturer’s instructions. DNase treatment was performed for 30 minutes at 37°C and DNase was removed via phenol-chloroform extraction. rRNA-depleted RNA was used to generate the RNA-sequencing libraries using NEB Ultra II directional kit (New England Biolabs). Samples from Keratinocytes were multiplexed and sequenced on a HiSeq4000 PE 150 sequencing platform (Illumina). RNA from transfected FaDu cells was used to generate RNA-sequencing libraries using TruSeq stranded total RNA (Illumina). Samples were multiplexed and sequenced on a NovaSeq 6000 SP PE 150. RNA from SCC25 cells and clones was used to generate RNA-sequencing libraries using stranded Total RNA prep with Ribo-zero plus (Illumina). Samples were multiplexed and sequenced on a NovaSeq 6000 SP/S1 PE 150. RNA from SRSF2 knockout and corresponding control mice was used to generate RNA-sequencing libraries using TruSeq Stranded Total RNA Gold (Illumina). RNA from SRSF2^P95H^ and corresponding control mice was used to generate RNA-sequencing libraries using stranded Total RNA prep with Ribo-zero plus (Illumina). Samples were multiplexed and sequenced on a NovaSeq 6000 SP PE 150. Three to five replicates were sequenced per condition.

Cells were isolated from mouse back skin as previously described ^88^. Briefly back skin was shaved and fat tissue scraped off from the dermal side. Skin pieces were then floated in 0.25% trypsin for 2 hours at 37°C, shaking. Trypsin was neutralised with Calcium free DMEM (Life Technologies) supplemented with FBS and filtered through a 70 μm filter. Cells were pelleted and then washed before resuspending them in 1% BSA for flow sorting. TdTomato-positive cells were collected in TRIZOL (Thermo Fisher). RNA from tdTomato positive SRSF2 knockout and P95H mutant cells and reporter positive control cells was extracted using the Direct-zol RNA microprep kit (Zymo Research), following manufacturer’s instructions. RNA-sequencing libraries were prepared using stranded Total RNA prep with Ribo-zero plus (Illumina). Samples from knockout mice and controls were multiplexed and sequenced on a NovaSeq 6000 SP PE 150. Samples from mutant mice and controls were multiplexed and sequenced on the NextSeq2000. Three replicates per condition were sequenced.

For thiol(SH)-linked alkylation for the metabolic sequencing of RNA (SLAM seq) ^42^, 4-thiouridine (s^4^U) labelling and RNA isolation was performed according to manufacturer’s instructions (Lexogen). Briefly, s^4^U was added to the culture medium of FaDu cells at a concentration of 500 μM for 15, 30 or 60 minutes, after which cells were harvested by adding TRIZOL. RNA was extracted under reducing conditions in the dark and quantified by nanodrop (NanoDrop One, Thermo Scientific). 5 μg of RNA was then treated with iodoacetamide and incubated at 50°C for 15 minutes. RNA was precipitated and libraries generated using the QuantSeq 3’ mRNA sequencing library prep kit (Lexogen) according to the manufacturer’s instructions. All samples were multiplexed and sequenced on a NovaSeq 6000 SR 100 SP sequencing platform (Illumina). Three to five replicates were sequenced for control and SRSF2 depleted samples per time point.

Cut&Run experiments were performed according to manufacturer’s instructions (Cell Signalling Technologies), with minor adjustments. Cells were harvested at room temperature and counted using the Countess II automated cell counter (Invitrogen). 100,000 cells were used per reaction. Initially cells were bound to activated concanavalin A magnetic beads. Permeabilization and incubation with primary antibody was then performed overnight at 4°C. pAG-MNase enzyme binding was then carried out for 1 hour at 4°C. DNA digestion was activated by adding calcium chloride to the reaction mixture and carried out at 4°C for 30 minutes. The reaction was then stopped and DNA diffusion was done by incubating samples at 37°C for 30 minutes. DNA purification was then carried out using DNA purification buffers and spin columns (Cell Signalling Technologies).

### RNA sequencing analyses

RNA sequencing data were processed by the DKFZ One Touch Pipeline (OTP) ^89^ using the RNA-seq workflow version 1.3.0 (https://github.com/DKFZ-ODCF/RNAseqWorkflow) in combination with the workflow management system Roddy version 3.5.8 or 3.5.9 (https://github.com/TheRoddyWMS/Roddy). In brief, data were aligned against the appropriate reference genome: 1KGRef_PhiX (generated from the 1000 Genomes assembly, based on hs37d5 and including decoy sequences merged with PhiX contigs to be able to align spike in reads) or GRCm38mm10_PhiX (based on GRCm38 merged with PhiX contigs to be able to align spike in reads) using the STAR aligner version 2.5.3a ^90^. Duplicate marking was performed using Sambamba version 0.6.5 ^91^ and quality control was performed using Samtools version 1.6 flagstat ^92^ as well as rnaseqc version 1.1.8 ^93^. FeatureCounts of the Subread package version 1.5.1 ^94^ was used for gene specific quantification of reads on the appropriate gene annotation: GENCODE version 19 or GENCODE version M12 in the strand-specific counting mode.

Raw gene count values were then used as input for a differential gene expression analysis using DESeq2 version 1.28.1 ^95^ within R version 4.0.0 R Core Team (2020). R: A language and environment for statistical computing. Foundation for Statistical Computing, Vienna, Austria. https://www.R-project.org/), which was made using default conditions besides fitType=“local” for each experiment separately. Results tables were generated for each treatment compared to the appropriate control.

For the detection of unique gene fusions Arriba version 0.8 ^96^ was used. The fusions listed in the output file with all fusions that pass all of Arriba’s filters was further filtered to only contain high confidence fusions and from these unique pairs of genes were counted.

### SLAM sequencing analyses

Adapter sequences were removed and only reads with a minimum length of 21 were kept using cutadapt version 3.4 (https://cutadapt.readthedocs.io/en/stable/) with the following parameters: -a “AGATCGGAAGAGCACACGTCTGAACTCCAGTCAC” and -m 21. The “all” option of the SlamDunk package version 0.4.3 ^42,97^ was used to align sequencing reads and perform read counting of total transcripts as well as newly transcribed RNAs, using the following parameters: −5 12, -n 100, -c 1, -m, -rl 101, --skip-sam, as reference GRCh38.dna.primary_assembly was used, in combination with a bed file based on Ensembl Genes version 104 that was modified by in-house scripts in conjunction with bedtools merge version 2.29.2 ^98^ to only include unique, non-overlapping 3’UTRs per gene that have a gene name assigned.

The count values of ReadCounts (total mRNAs) and TcReadCounts (newly-transcribed mRNAs) were then summed up per gene for each sample. These were used as input for a differential expression analysis using DESeq2 version 1.28.1 ^95^ within R version 4.0.0 (R Core Team (2020). R: A language and environment for statistical computing. Foundation for Statistical Computing, Vienna, Austria. https://www.R-project.org/). Both categories (total and newly-transcribed genes) were considered separately, meaning that a DESeqDataSet was set up for each of the two categories. The differential expression analysis was then made using default conditions besides fitType=“local”. Results tables were then generated for each time point and for each category (total and newly-transcribed) separately. Results were then used to assess gene expression in a quantitative manner using density plots and cumulative frequency plots, as well as gene ontology analysis.

### Quantifying bi-directional transcription

The distance to the closest neighbour of protein coding genes in any orientation was identified as described in ^49^. Protein coding genes were first extracted from the gencode version 38 annotation file and these were then used as input for bedtools version 2.24.0 closest ^98^ command with varying parameters depending on the direction.

How distance and fold change after SRSF2 depletion correlate was visualised via violin plots ^49^. The proportion of genes with a neighbour within 1000 base pairs when categorised by fold change, greater than 0.3/less than −0.3, as determined by DESeq2.

### Cut and Run analyses

FastQ files were processed using the nf-core CUT&RUN pipeline ^99^. Briefly, adaptors were trimmed using TrimGalore ^100^ and reads were aligned to the human reference genome Hg19 (targets), or to the yeast reference genome R64-1-1 (spike-in) using Bowtie2 ^101^. Bedtools were used to generate bedGraph files and bedGraphToBigWig was used to produce BigWig files.

Differentially expressed genes were identified from SLAM-seq data using DESeq2 ^95^, genes were filtered to have a mean FPKM > 5 and an adjusted p-value of 0.05. For each gene its “Ensembl Canonical” transcript was selected. The number of reads aligned to a 500 bp window centred on the transcription start site (TSS) and the number of reads aligned within the gene body (from transcript start −250bp to transcript end + 250bp) were counted. A pausing index was then computed as the ratio of reads at the TSS divided by the number of reads in the gene body.

Two methods were used to compute significance of the pausing index between knock down and control. First, a t-test was performed per gene for knock down replicates and control replicates. These p-values were then combined using Fisher’s method. Second, the log ratio of the mean TSS enrichment for knock down replicates was divided by the log ratio of the mean TSS enrichment for the controls and a t-test was then performed on these log ratios.

### Splicing analyses

Differential splicing analysis with LeafCutter (v0.2.7) and Majiq (v2.2) was carried out using the Baltica Framework ^102–104^ using default parameters. In brief, LeafCutter differential splicing used a minimum samples per group of two and a minimum samples per intron of two. For Majiq Voila tsv the threshold was set to 0.1. Individual introns were examined and significant events were defined as those with an adjusted p-value of 0.05 (Leafcutter) and a probability of non-changing greater than 0.95 (Majiq). Alternative splicing (AS) events were calculated as percent spliced in (PSI) representing the proportion of splice site usage for a particular AS event per experimental group and indicates effect size ^105^. The deltaPSI was calculated as the difference of two PSI values across two experimental groups. Gene ontology for genes with differential exon inclusion was conducted with GOrilla ^106,107^

### SRSF2 transcriptome binding

SRSF2 motifs were obtained from the oRNAment database ^108^. These were then filtered to canonical chromosomes. CLIP-seq peaks were downloaded and processed from the Dorina database and aligned to the Hg38 assembly using the UCSC chain files ^109^. Peak enrichment analysis was then performed using ChIPseeker (v1.33.1), org.Hs.eg.db (v3.14.0) and the intersection of the SRSF2 CLIP-seq experiments and SRSF2 motifs as input ^110,111^. The intersection totalled 4421 peaks, of which 4417 were annotated by ChIPseeker. 30% of peaks were annotated to promoter regions, 2% to the 5’UTR and 64% to other exons. Promoters were defined as the 6 kb around the transcription start site. First exons were defined as the target for enrichment analysis and the ChIPseeker::plotAvgProf function with 50 bins and 1000 resampling steps were parameters used to obtain enrichment profiles.

Human CLIP-seq data for wild-type and P95L mutant SRSF2 (Figure 6A) were obtained from ^112^. Bed files were downloaded and analysed with ChIPseeker (v1.37.0). Peak profiling was executed with the plotPeakProf function to transcript bodies, using a relative distance from −20% TSS to 20% the TES. 95% confidence interval was obtained via bootstrapping and the number of bins set to 800. The peak score was taken in account.

### Sample sizing and collection

No statistical methods were used to predetermine sample size, but a minimum of three samples were used per experimental group and condition. The number of samples is represented in the graphs as one dot per sample. Samples and experimental animals were randomly assigned to experimental groups. Sample collection was also assigned randomly. Sample collection and data analysis were performed blindly whenever possible. Whenever possible automated quantifications were performed using the appropriate software.

## Data availability

Mouse RNA sequencing data are available on GEO (GSE221444). Human sequencing data are available on GEO (GSE222021). Results are in part based on published mouse CLIP-seq data (GSE44591) and human CLIP-seq data (GSE164666). CLIP-seq peaks were downloaded and processed from the Dorina database and aligned to the Hg38 assembly using the UCSC chain files.

## Code availability

CUT&RUN analyses: https://github.com/CompEpigen/cutandrun_SRSF2_Wagner2023. Splicing analyses: https://github.com/dieterich-lab/Baltica.

## Acknowledgments

We thank all DKFZ core facilities for their support, and in particular the light microscopy, flow cytometry, and the genomics and proteomics facilities. Special thanks also go to the Omics IT and data management core facility (ODCF), and all members of staff of the DKFZ Central Animal Laboratory. We also thank the genomics core facility at the Cancer Research UK Cambridge Institute for help with some sequencing and data transfer. This work was funded by the Helmholtz Association (W2/W3-106) and Cancer Research UK (CR-UK; C10701/A15181). Some figure panels were created with BioRender.com.

## Author contributions

REW, MF: Designed experiments and wrote manuscript

REW, LA, AB, SD: Performed experiments and analysed data

DS: Collected and processed tissue samples.

TB-B, AH-M, ES, JP: Performed computational analyses

SB: Established and performed computational analyses

DTO, CP, CD, PL: Supervised computational analyses

PS: Supervised comet assay experiments and data analysis

JP: Performed some of the sequencing, data visualization and statistical analyses.

NM-D: Performed DMBA/TPA experiments

## Declaration of interests

All authors declare no conflict of interest

---

**Supplementary Figure 1.**
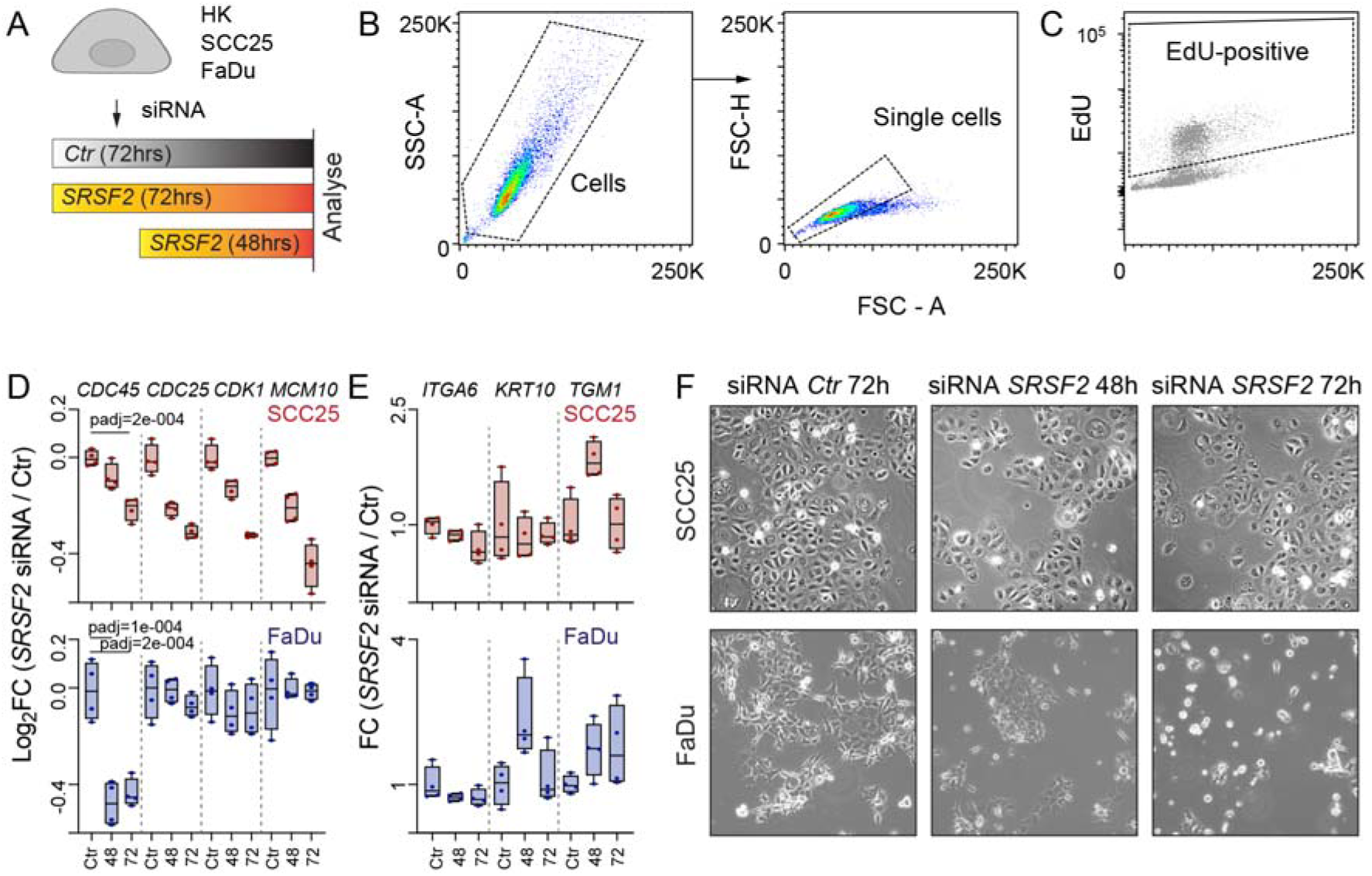
Depletion of SRSF2 causes cell cycle arrest. (**A**) Schematic overview of transfection experiments using a scrambled control (Ctr) and *SRSF2* siRNAs in primary human keratinocytes (HK) and the squamous cell carcinoma lines SCC25 and FaDu. Cells were analysed 48 or 72 hours (h) after transfection. (**B,C**) Flow cytometry gating (B) to quantify EdU-positive cells (C). (**D**) Log_2_ fold change (FC) of cell division cycle 45 (*CDC45*), cell division cycle 25 (*CDC25*), cyclin dependent kinase 1 (*CDK1*) and minichromosome maintenance 10 (*MCM10*) RNA levels in SCC25 (top) and FaDu (bottom) in Ctr and SRSF2-depleted conditions. (**E**) FC of Integrin α6 (*ITG*α*6*), keratin 10 (*KRT10*) and transglutaminase 1 (*TGM1*) RNA levels in SCC25 (top) and FaDu (bottom) in Ctr, and SRSF2-depleted conditions. N = 4 transfections (D,E). (**F**) Bright field images of Ctr and SRSF2-depleted SCC25 (top) and FaDu (bottom) cells 48 and 72 h after transfection. Box plots show minimum, first quartile, median, third quartile, and maximum (D,E). Dunnett’s multiple comparisons test (D). Exact p-values are indicated.

**Supplementary Figure 2.**
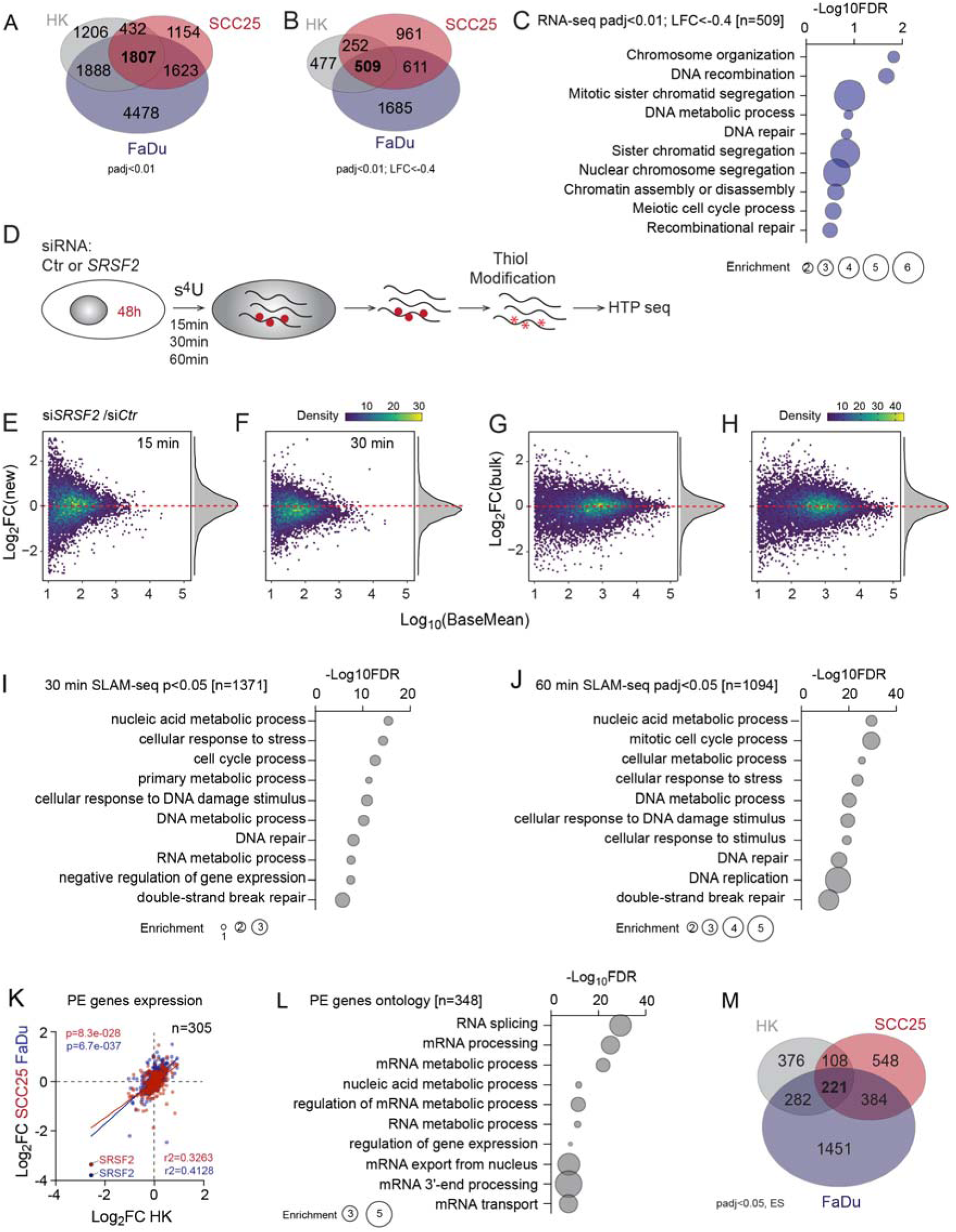
Splicing changes are inconsistent across cell lines after knockdown of SRSF2. (**A**) Venn diagram showing significantly (padj < 0.01) differently expressed genes in *SRSF2*-depleted HK, SCC25 and FaDu cells. (**B**) Illustration of experimental set-up for transcript metabolic labelling using 4 thiouridine (s4U) followed by RNA sequencing. (**C-F**) Change of global nascent (C, D) and bulk (E, F) RNA levels after 15 (C,E), 30 (D, F) minutes (min) after 4sU labelling in si*SRSF2* versus si*Ctr* cells. Shown are all genes with a base mean >10. (**G, H**) Gene ontology analysis using significantly (padj<0.05) differentially newly transcribed genes 30 (G) or 60 (H) minutes after metabolic labelling (n = genes). (**I**) Correlation of expression changes of genes containing an ultra-conserved poison exon (PE) cassette in *SRSF2*-depleted HK, SCC25 and FaDu cells (n = genes). (**J**) Gene ontology analysis of genes containing ultra-conserved PEs. (**K**) Venn diagram showing genes with significant exon skipping events (padj < 0.05) in *SRSF2*-depleted HK, SCC25 and FaDu cells.

**Supplementary Figure 3.**
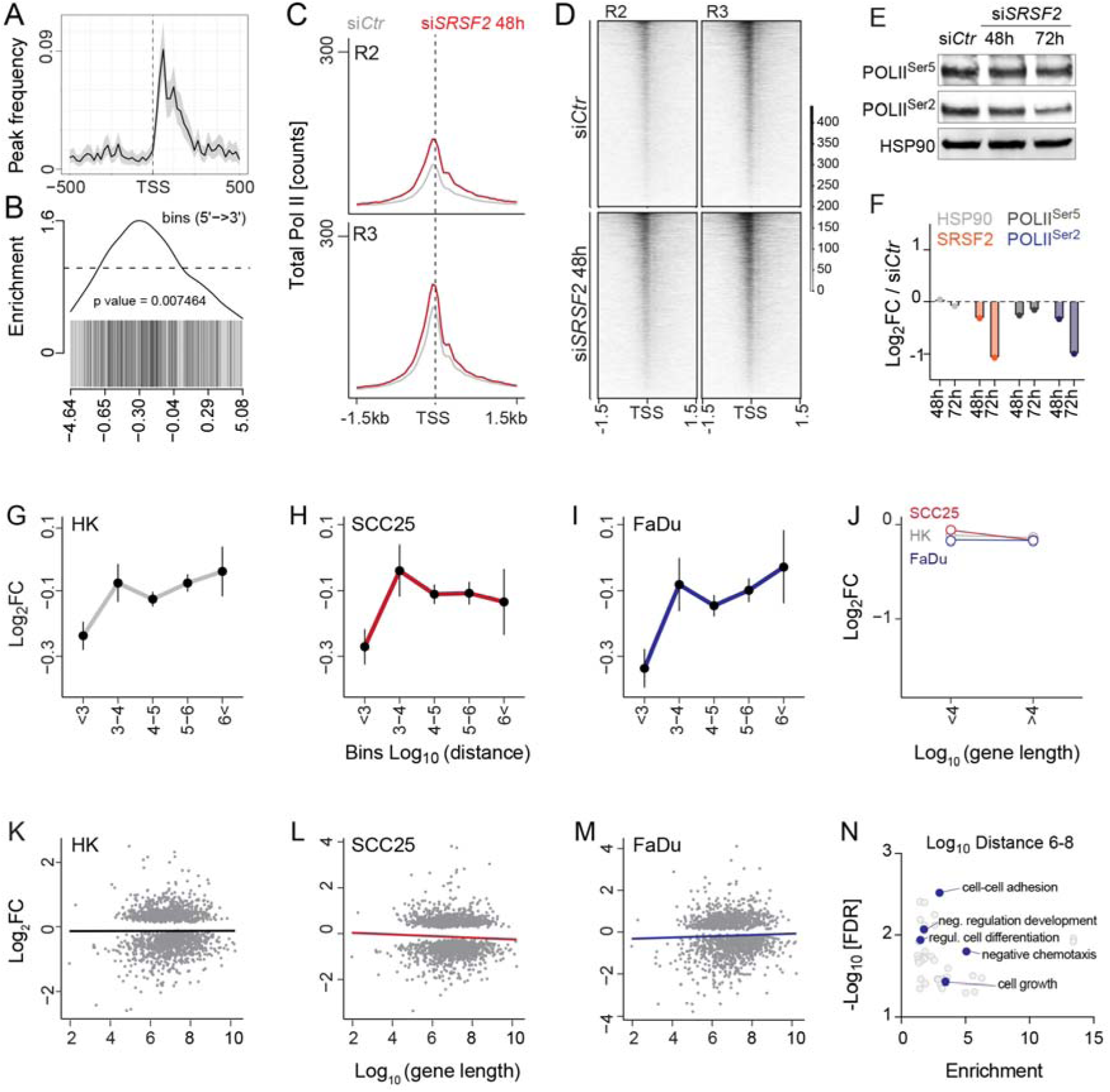
SRSF2 is required for efficient transcription of a subset of genes. (**A**) Frequency of SRSF2-binding sites 500 bp up- and down-stream the transcriptional start sites (TSS) in first exons. (**B**) Enrichment of SRSF2-binding sites in the first exon of genes significantly (padj < 0.05) differently transcribed in SLAM-seq experiments (60 minutes). P = Fisher’s exact test. (**C, D**) Total Pol II occupancy (C) and heatmaps (D) at transcription start sites (TSS) in si*Ctr* and si*SRSF2* cells 48 hours (h) after transfection. Shown are two out of three replicates. R2: Replicate 2, R3: Replicate 3. (**E,F**) Western Blot (E) and quantification (F) of phosphorylated Pol II at serine 2 and 5 (Pol II^Ser^^2^, Pol II^Ser^^5^) in control (si*Ctr*) and SRSF2-depleted (si*SRSF2*) FaDu cells 48 or 72 h after transfection. (**G-I**) Common differentially expressed (p < 0.01) genes in primary human keratinocytes (HK) (G) and the squamous cell carcinoma lines SCC25 (H) and FaDu cells (I) (n = 1807) separated by their distance to the next up-stream protein coding genes in anti-sense direction. Shown is mean ± SEM. (**J**) Log_2_ fold change (FC) of bi-directionally transcribed genes smaller than 10kb (< 4) or longer than 10 kb away (> 4) in HK, SCC25 and FaDu cells. P = Wilcoxon rank sum test with continuity correction. (**K-M**) Common differentially expressed (p<0.01) genes in HK (K), SCC25 (L) and FaDu (M) cells separated by their length. (**N**) Gene ontology of anti-sense transcribed genes with a distance between 1 and 100 Mb from each other.

**Supplementary Figure 4.**
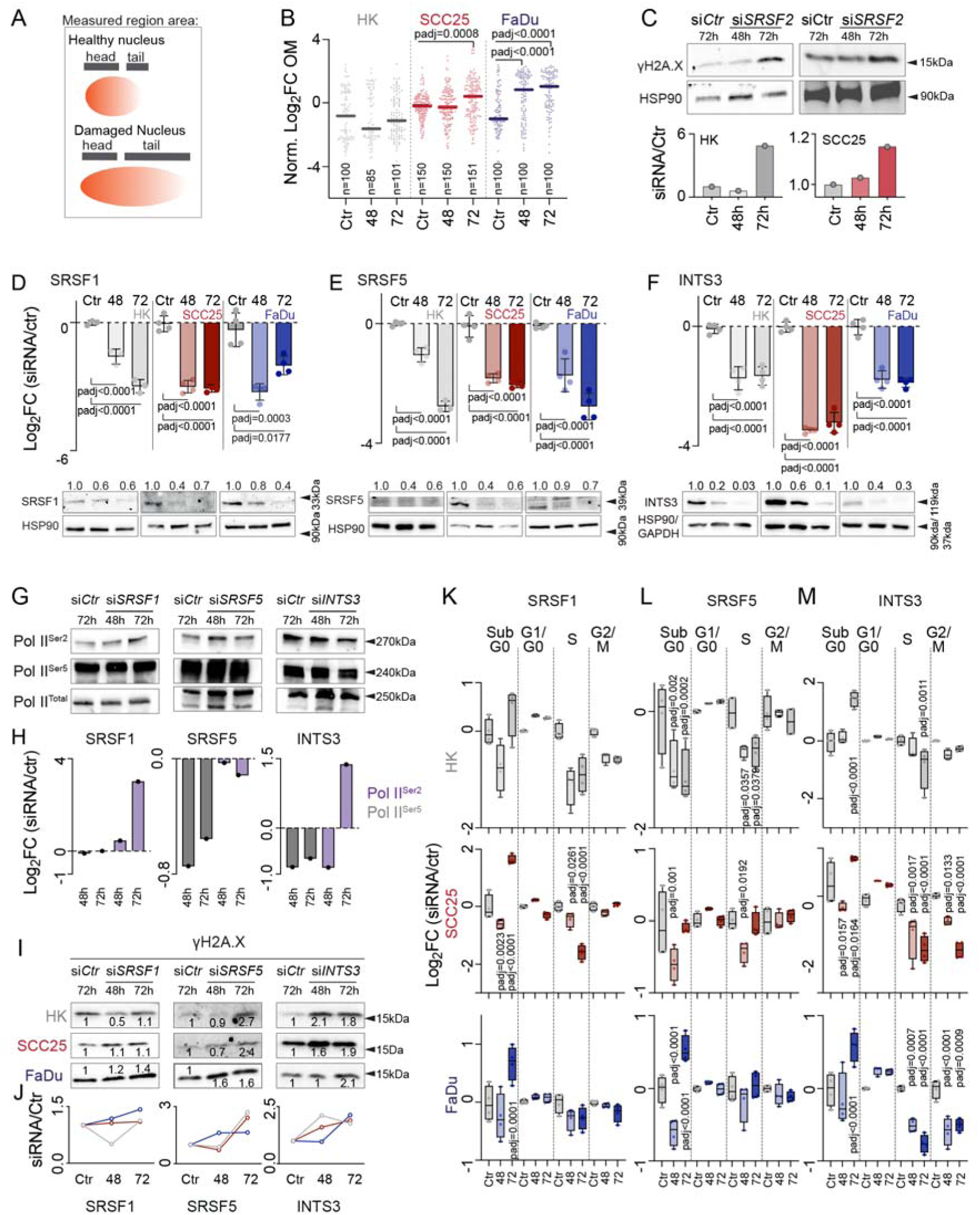
Depletion of other SR proteins does not lead to increased DNA breaks. (**A**) Illustration of comet assay measurements. (**B**) Log_2_ fold change (FC) of the normalized (norm.) olive tail moment (OM) relative to scramble control (Ctr) siRNA in primary human keratinocytes (HK) and the squamous cell carcinoma lines SCC25 and FaDu 48 or 72 hours (h) after transfection of *SRSF2* siRNA (n = number of cells). **(C**) Western Blot (upper panels) and quantification (lower panels) of phosphorylated γH2AX protein levels by measuring band intensity in si*Ctr* and si*SRSF2* transfected HK (left) and SCC25 (right) cells after 48 or 72h. HSP90: Loading control. (**D-F**) Log_2_ FC RNA levels (top panels) and protein levels (bottom panels) of SRSF1 (D), SRSF5 (E) and INTS3 (F) in HK (gray), SCC25 (red) and FaDu (blue) cells transfected with Ctr siRNAs for 72h or *SRSF1, 5 or INTS3* siRNAs for 48h or 72h (n = number of transfections averaged over 3 technical replicates). HSP90 or GAPDH: Loading control. (**G, H**) Western Blot (G) and quantification (H) of phosphorylated Pol II at serine 2 and 5 (Pol II^Ser^^2^, Pol II^Ser^^5^) in si*Ctr* or *SRSF1-*, *SRSF5-* and *INTS3*-depleted HK cells 48 or 72h after transfection. (**I, J**) Western Blot (G) and quantification (H) of γH2AX protein levels in si*Ctr* and si*SRSF1*, *siSRSF5-* and *siINTS3* transfected HK, SCC25 and FaDu cells for 48 or 72h. Loading controls for (G-J) are HSP90 or GAPDH shown in (D-F). (**I-K**) Number of *SRSF1* (K), *SRSF5* (L) *and INTS3* (M) -depleted HK (top), SCC25 (middle) and FaDu (bottom) cells in the different phases of the cell cycle 48 and 72h after transfection. Data are relative to Ctr (72h) (n = number of transfections). Shown is mean (B) or mean± SD (D-F). Box plots show minimum, first quartile, median, third quartile, and maximum (K-M). Dunnett’s multiple comparisons test (D-F). Exact p-values are indicated.

**Supplementary Figure 5.**
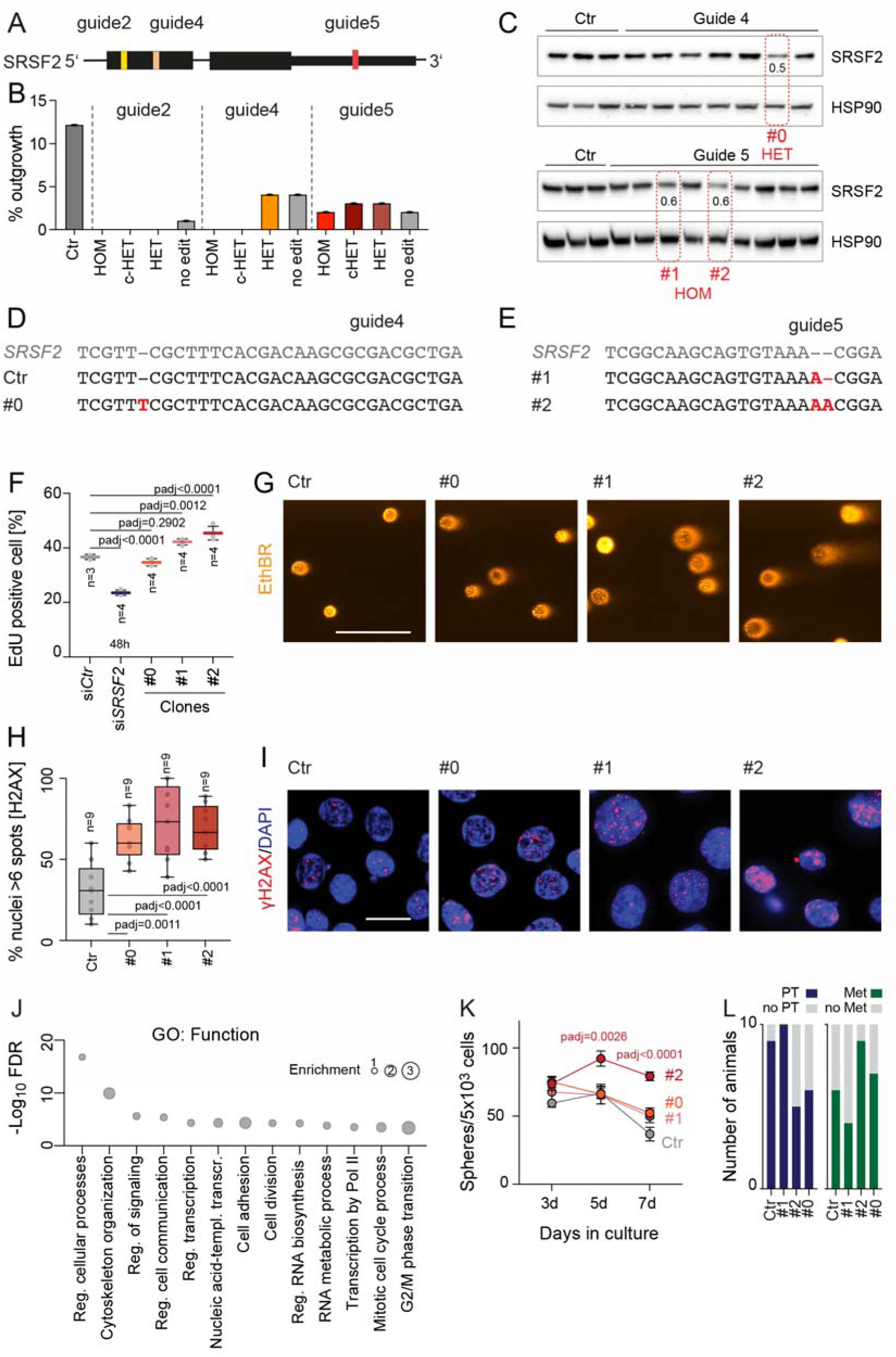
A two-fold reduction of SRSF2 levels causes DNA damage without affecting cell cycle. (**A, B**) Illustration of CRISPR/Cas9 approach (A) and clonal outgrowth (B) targeting SRSF2 with three different guide RNAs (guide2, 4 and 5). (**C-E**) SRSF2 protein levels (C) and targeted sequence (D,E) in control (Ctr) or SRSF2 edited clones using guide 4 and 5 resulting in homozygous (HOM; #0) or heterozygous (HET; #1 and #2) mutations in *SRSF2*. Number in (C): band intensity of SRSF2 (Fiji). HSP90: loading control. Red nucleotides (D,E): mutations in *SRSF2*. (**F**) Percentage of EdU positive Ctr and SRSF2 knockdown clones #0,#1 and #2 (n = number of EdU incubations; each point is average of 10,000 events). (**G**) Representative Comet assay images showing ethidium bromide (EthBR) stained Ctr and #0, #1 and #2 cells (n = 4 transfections averaged over 3 technical replicates). (**H, I**) Quantification of nuclei with more than six γH2AX foci (H) and representative fluorescence stainings for γH2AX (I) in cells from Ctr and #0, #1 and #2 clones (n = 9 images). (**J**) Gene ontology analysis of commonly significantly (padj<0.01) up-regulated (Log2FC>0) genes in all clones. (**K**) Growth of tumour spheres for 3, 5, and 7 days (d) formed by Ctr or #0, #1 and #2 clones (n = 6 tumour sphere assays). (**L**) Quantification of total number of primary tumour (PT) and lymph node metastases (Mets) formed by Ctr and #0, #1 and #2 clones at day 27 (n = 10 mice per clone). Shown is mean± SD (F). Box plots show minimum, first quartile, median, third quartile, and maximum (H). Dunnett’s multiple comparisons test (F,H). Exact p-values are indicated.

**Supplementary Figure 6.**
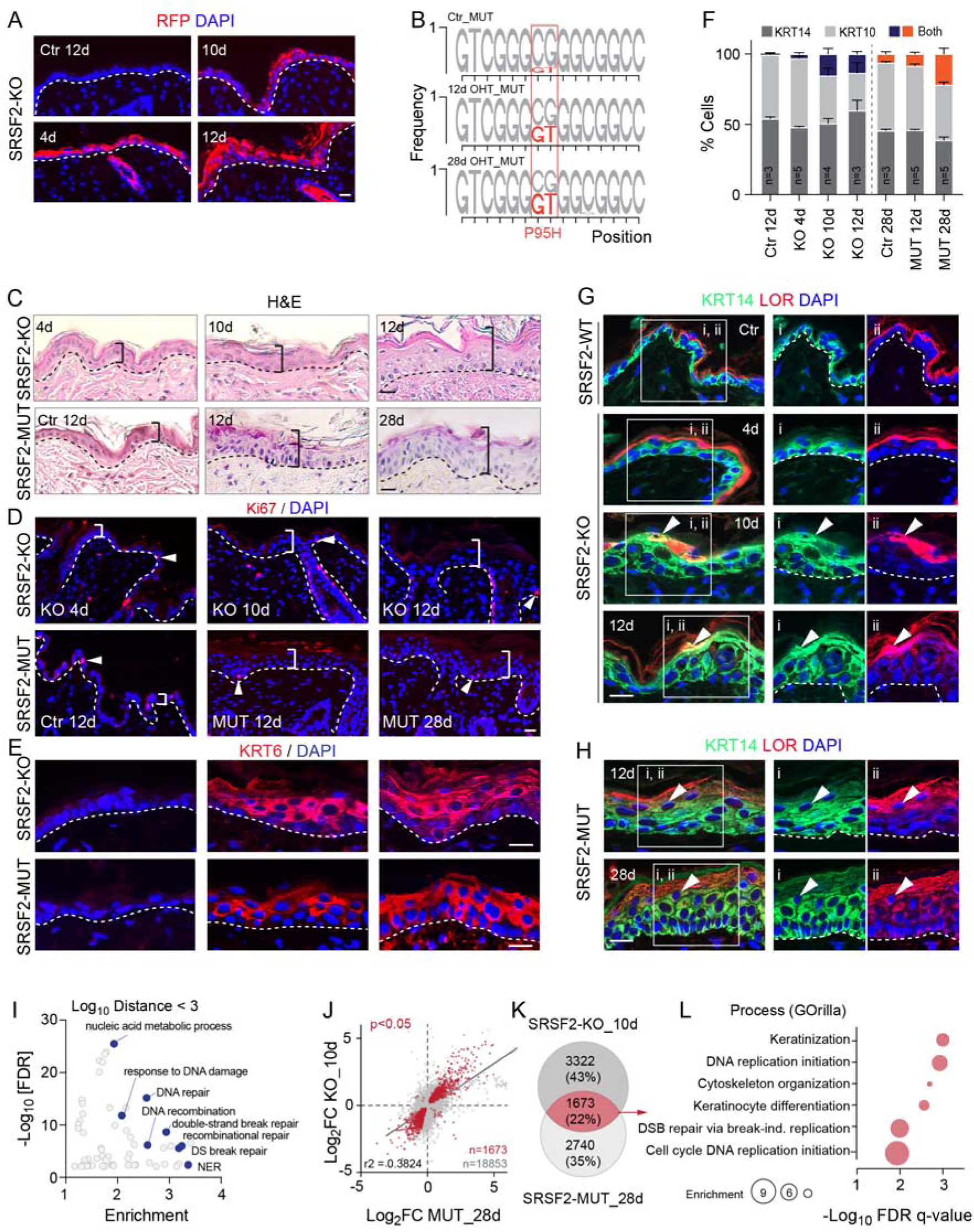
Loss of SRSF2 impairs differentiation and triggers tissue repair response in mouse epidermis. (**A**) Immunofluorescence staining for tdTomato expression using a Red Fluorescent Protein (RFP) antibody in mouse back skin treated with 4-hydroxytamoxifen (OHT) for 4, 10, or 12 days (d). (**B**) Frequency of the proline-to-histidine (P95H) mutation (MUT) in *Srsf2* transcripts in control (Ctr) and mutated (SRSF2-MUT) back skin after 12d and 28d OHT treatment. Orange: Nucleotide substitutions (P95H). (**C**) Mouse skin histology (hematoxylin and eosin) of OHT-treated *Srsf2* knockout (SRSF2-KO) and SRSF2-MUT epidermis. Brackets: Epidermal thickness. Dashed lines: Basement membrane. (**D**) Immunofluorescence staining for Ki67 in SRSF2-KO and -MUT mouse epidermis after the indicated OHT treatment. Brackets: Epidermis thickness. Arrowheads: Ki67-positive cells. Dashed lines: Basement membrane. (**E**) Immunofluorescence staining for keratin (KRT) 6 in back skin from SRSF2-KO and -MUT mice after the indicated time point of OHT-treatment. (**F**) Percentage (%) of cells staining KRT14 and KRT10 double-positive (Both) in SRSF2-KO -MUT epidermis (n = number of mice). (**G, H**) Immunofluorescence staining for KRT14 and loricrin (LOR) in back skin from SRSF2-KO (G) and -MUT (H) mice after the indicated time of OHT-treatment. White square: highlighted area shown in i and ii. Arrowheads: Double-positive cells. Dashed lines: Basement membrane. DAPI: nuclear counterstain (A, D, E, G, H). (**I**) Gene ontology analysis of bi-directionally transcribed mouse genes. (**J**) Correlation of differentially expressed genes 10 days after SRSF2 deletion (KO_10d) and 28 days of SRSF2 mutation (MUT_28d). Red: Significant (p<0.05) genes. (**K**) Overlap of significantly (p<0.05) differently expressed genes in KO_10d and MUT_28d epidermis. (**L**) Gene Ontology (GO) analyses using overlap of genes (red) in (K) (GOrilla). Background: All expressed genes. Shown is mean± SD (F).

**Supplementary Figure 7.**
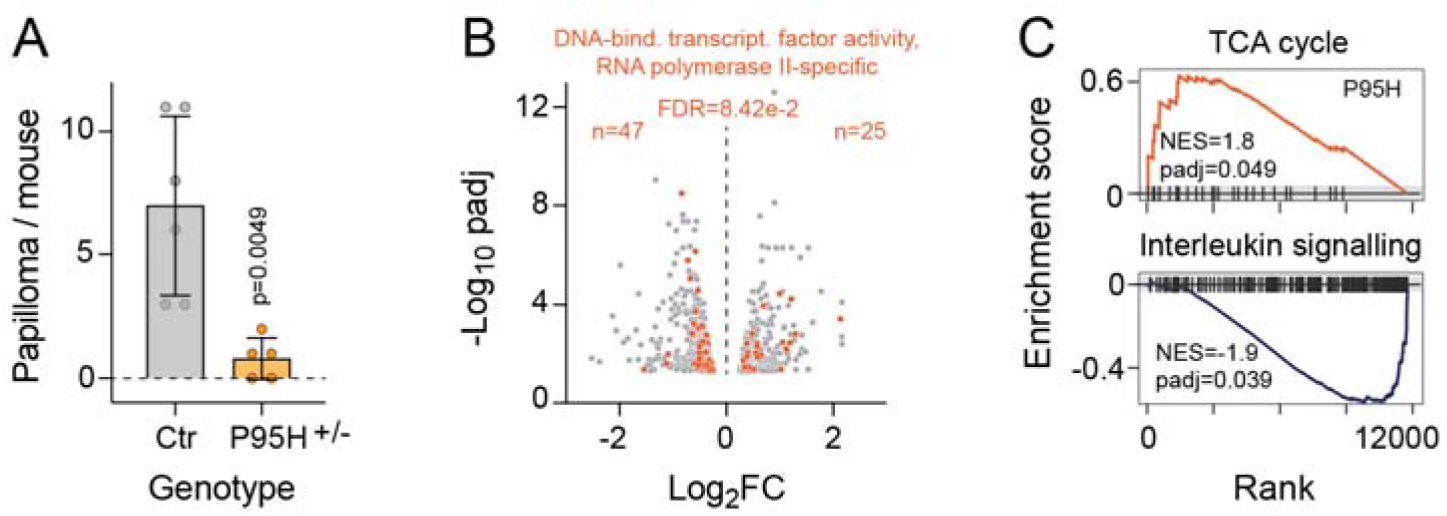
Srsf2 P95H+/− epidermal cells are metabolically different. (**A**) Number of papillomas per mouse in control (Ctr) and *Srsf2* P95H+/− skin after treatment with carcinogens. Shown is mean± SD. Dunnett’s multiple comparisons test (F,H). Exact p-values are indicated. (**B**) Volcano plot showing significantly (padj<0.01) de-regulated genes in P95H+/− compared to ctr tdTomato-positive cells. Orange: Genes of the GO: 0000981 (FDR=8.42e-2). (**C**) Significantly enriched gene sets in SRSF2-P95H+/− cells. NES: normalized enrichment score.

## Notes

### Competing Interest Statement

The authors have declared no competing interest.

